# Convergent latitudinal erosion of circadian systems in a rapidly diversifying order of fishes

**DOI:** 10.1101/2025.05.28.656707

**Authors:** Daniel B. Wright, Yanfan Zhang, Jacob M. Daane

## Abstract

Biological clocks enable organisms to anticipate cyclical environmental changes. Some habitats, such as those at high latitudes or deep sea, experience seasonally diminished or absent diel cues upon which species entrain their circadian rhythms. Fishes of the order Perciformes have rapidly diversified and adapted to these arrhythmic ecosystems, raising the possibility that evolutionary modifications to their circadian biology contributes to their success as one of the most species-rich orders of vertebrates. Here, we used a comparative genomic approach to investigate patterns of biological clock gene loss and circadian rhythms across 33 perciform and six outgroup species. We found both widespread and lineage-specific loss and relaxed selection in core clock genes, particularly in the convergently evolving polar and deep-sea Notothenioidei and Cottioidei suborders. This trend of circadian gene loss was significantly correlated with latitude, with higher-latitude species showing greater loss. Whether these losses and relaxed selection lead to changes in circadian rhythms is unknown for most perciforms. To address this, we performed metabolic phenotyping on three notothenioid species and found no circadian metabolic oscillations during the late austral fall, including in the sub-Antarctic *Eleginops maclovinus*, sister to the Antarctic adaptive radiation. We propose that diminished reliance on endogenous biological clocks may be an adaptive feature that facilitates the survival and diversification of perciform fishes in polar and arrhythmic environments.

## Introduction

Most habitats on Earth experience regular environmental cycles, driven by celestial rhythms such as Earth’s rotation and orbit, which produce predictable patterns of light and darkness, and by lunar forces that govern tidal movements. In response to these rhythms, species have evolved internal molecular biological clocks that enable them to anticipate the optimal time to invest energy in activities such as foraging in sync with the cycle of their ecosystem (1). These endogenous circadian rhythms have been observed across the tree of life, including bacteria, archaea, plants, and metazoans (2, 3) and at every stage of an organism’s life cycle (4, 5). Biological clocks can be calibrated, or “entrained”, to external cues. The most common environmental cue, also known as a *zeitgeber* (time giver), is light, though food availability, lunar cycles, temperature, and tidal forces have all been reported to influence circadian periodicity (6–8). Many genes are regulated by the biological clock; in mammals, up to 43% of all protein-coding genes exhibit circadian expression patterns (9). The ubiquitous presence of circadian rhythms across biota highlights their central importance in organismal fitness, and disruption to circadian cues can lead to elevated disease risk (10–13).

Many species have evolved in environments where common environmental zeitgebers are absent or seasonally disrupted and yet thrive in these habitats (14). High-latitude polar ecosystems experience extreme seasonal variation in periodic cues such as the light/dark cycle that typically entrain circadian rhythms (15). These extreme shifts in diel periodicity are considered a barrier that challenges poleward-migrating species that otherwise depend on regular circadian rhythms (16). Polar vertebrates exhibit a spectrum of adaptations to extreme high-latitude light/dark cycles, including sustained circadian entrainment through polar summers and winters, seasonal arrhythmias, and ultradian or free-running rhythms often driven by non-photic cues (15).

The large order Perciformes (*sensu stricto*), which contains about 9% of all teleost fish species (∼3,300) (17), is overrepresented in high-latitude environments, comprising ∼66% of high-latitude fish diversity and encompassing four of the most rapidly diversifying marine clades: Cryonotothenioidei, Sebastidae, Zoarcidae, and Liparidae (18). These perciform radiations often involve diversification along a depth axis (19), with deep-sea invasions in Perciformes also correlated with latitude (20). Depth adaptations in this group are extreme; camera traps and trawl surveys have identified perciforms among the most abundant fishes at hadal (> 6,000 meter) depths (21–24). These deep invasions are attributed in part to reduced stratification of temperature and pressure at high latitudes, which lowers physiological barriers to depth invasions (20). However, another shared but underappreciated feature of these environments is the seasonal or permanent disruption of common circadian cues, particularly the light/dark cycle and feeding schedules.

Is there a unique genetic or physiological potential that underlies the successful and rapid diversification of Perciformes in extreme environments? Recent high-quality genome assemblies of notothenioids, hadal snailfishes, and deep-sea eelpouts separately uncovered loss of core circadian genes (25–28). These losses may reflect relaxed selection on biological clock genes in arrhythmic environments, but functional evidence remains limited, and data on circadian rhythms in perciforms are sparse. In field studies of geotagged Antarctic bullhead notothen *(Notothenia coriiceps)* along the Antarctic peninsula, adults showed no daily movement patterns during polar summer or winter, though activity, heart rate, and metabolism varied by season, suggesting possible circannual rhythms (29). In three-spined sticklebacks (*Gasterosteus aculeatus*) from Lake Témiscouata, weak and variable circadian locomotor activity was observed under light/dark cycles, with 82% of individuals arrhythmic in constant darkness (30). Notably, no diel expression of core clock genes (e.g., *arntl1a*, *clock1b*, *clock2*, *per1b*, *cry1b*) was detected, even under regular light/dark cycles (30). Similarly, two Arctic freshwater sculpins (*Cottus gobio* and *C. poecilopus*) showed seasonally shifting diurnal and nocturnal activity patterns, with *C. poecilopus* arrhythmic in summer (31), a pattern also seen in the marine shorthorn sculpin (*Myoxocephalus scorpius*) (32). These findings, collected around distinct measures of circadian rhythms, suggest that flexible or reduced circadian control may be a defining feature of this group.

It remains unclear whether disruptions to biological gene and circadian physiological rhythms are restricted to certain lineages or if they are widespread across Perciformes, which could suggest an ancestrally reduced reliance on circadian regulation that facilitates their expansion into arrhythmic habitats. We hypothesized that evolutionary plasticity in circadian rhythms and relaxed selection on clock genes enables the rapid diversification of Perciformes in such environments. To test this hypothesis, we analyzed core circadian gene status in 33 perciform and six outgroup species and experimentally assessed circadian metabolic rhythms in three notothenioid species, including *Eleginops maclovinus*, the sister lineage to the Antarctic adaptive radiation of notothenioids. Our results demonstrate that: (1) circadian gene loss and relaxed selection are convergent features in the rapidly diversifying suborders Notothenioidei and Cottioidei, common in polar and deep-sea habitats; (2) gene loss correlates with latitude; and (3) circadian rhythmicity is absent even in non-polar species like *E. maclovinus*, suggesting that flexible circadian control may be a feature that enabled polar diversification in this rapidly diversifying clade.

## Results

### Curation of core biological clock components in Perciformes

In vertebrates, the circadian clock is regulated through a transcription-translation feedback loop (TTFL). At the start of this circuit, the transcription factor proteins Circadian Locomotor Output Cycles Kaput (Clock) and Aryl hydrocarbon receptor nuclear translocator-like (Arntl, also known as Bmal) form a heterodimer that binds to E-box promoter elements across the genome. The Clock-Arntl heterodimer facilitates the transcription of Cryptochrome (Cry) and Period (Per) proteins, which ultimately bind the Clock-Arntl heterodimer and inactivate the complex. This feedback loop takes approximately 24 hours to complete, providing the central mechanism for synchronizing the cellular activity of most organisms with the Earth’s rotation (33).

To assess patterns of biological clock evolution in Perciformes, we investigated a panel of 30 well-conserved fish clock-associated genes (**Fig. 1**, **Table S1**)(34). This set includes paralogs of the core positive transcription factors: the *clock* genes (*clocka*, *clockb*), the *clock* homolog *npas2*, and the *arntl*/*bmal* genes (*arntl1a*, *arntl1b*, *arntl2a*, *arntl2b*) (33, 35, 36). We also included the negative regulators of the feedback loop: *period* paralogs (*per1a*, *per1b*, *per2a*, *per2b*, *per3*) and *cryptochrome* paralogs (*cry1a*, *cry1b*, *cry2*, *cry3a*, *cry3b*) (33, 37–40). In addition, we analyzed *cry4*, a photopigment-like cryptochrome implicated in light input and peripheral clock regulation (41), along with *cry5* and *cry-dash*, two cryptochrome family members involved in DNA repair that are light sensitive but are not considered core components of the biological clock (42, 43).

**Fig. 1.**
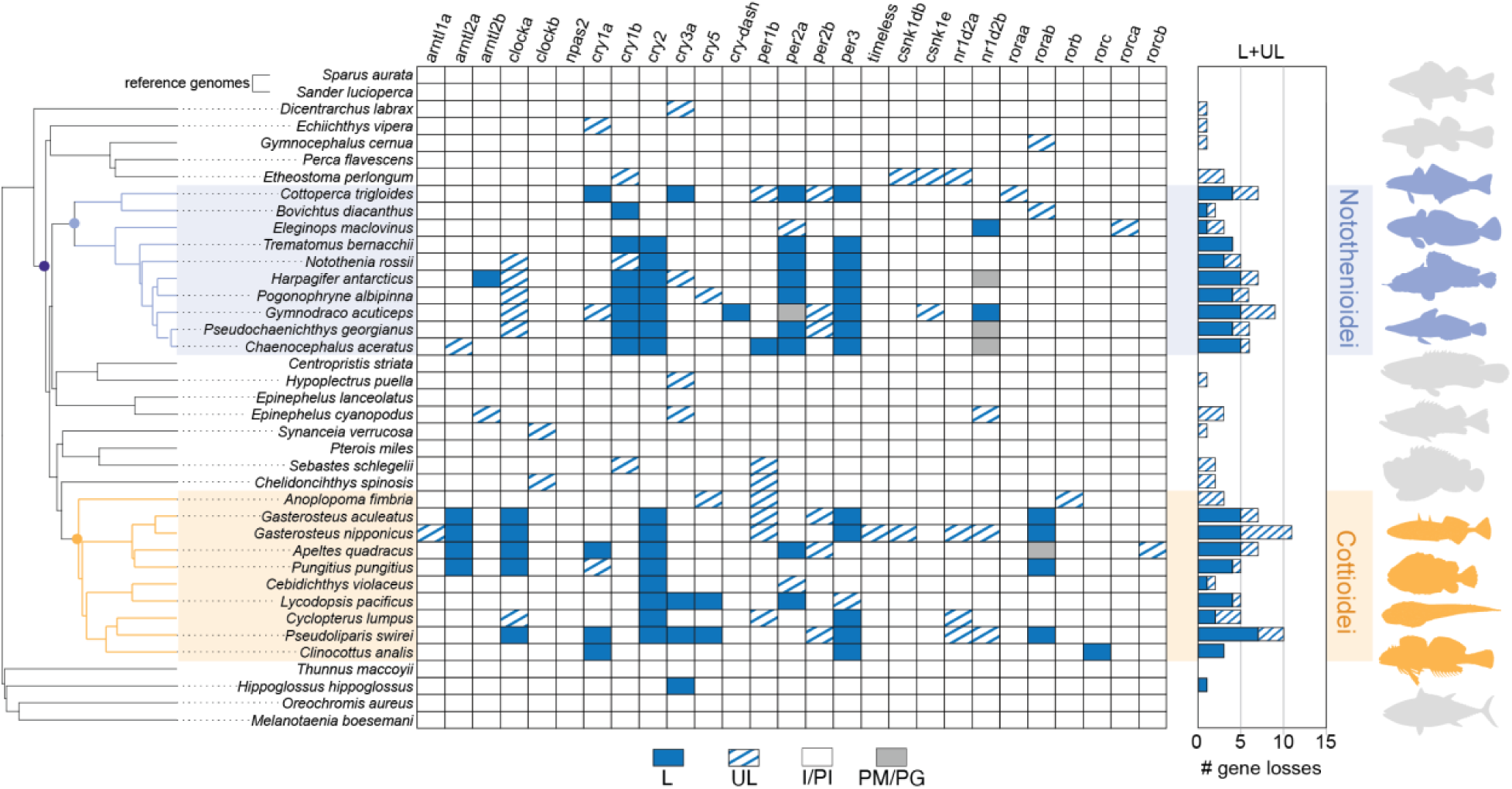
Loss of core biological clock genes in Perciformes. Phylogeny of Perciformes, including five outgroup species, pruned from Rabosky et al (18). Gene status was inferred from pairwise genome alignments between each species and one of two reference genomes, *Sparus aurata* or *Sander lucioperca*. For each gene in each species, the most complete annotation across both references was used. Losses (L; blue) indicate multiple truncating mutations within the central 80% of the coding sequence (CDS). Uncertain losses (UL; blue hash) reflect a single truncating or deletion variant within the middle 80% of the gene. Gray symbols denote genes that are partially missing (PM) or part of a paralogous group (PG), suggesting possible but highly uncertain gene loss or assembly artifacts. White indicates intact or partially intact genes (PI/I; white). Two clades with elevated gene loss are highlighted: Notothenioidei (blue) and Cottioidei (orange).

We further included regulatory proteins involved in the post-translational modification of the central Clock/Arntl and Cry/Per feedback loop. Among these are the Casein kinases (*csnk1da*, *csnk1db*, *csnk1e*), which phosphorylate Period proteins to regulate their stability, degradation, and nuclear localization (44–46). We also examined Timeless, a protein essential for clock function in *Drosophila* where it stabilizes Period proteins in the cytoplasm (47). Although its role in the vertebrate clock is less well defined, Timeless appears to have circadian functions and may link the clock to other cellular processes, such as the DNA damage response or interactions with cryptochromes (47).

Lastly, we included nuclear receptors that help stabilize circadian rhythms through transcriptional regulation. These consist of Nr1d family members (*nr1d1* [Rev-Erbα], *nr1d2a*, *nr1d2b* [Rev-Erbβ paralogs]), which repress the transcription of *arntl* (Bmal) and *clock/npas2* in a rhythmic manner (48), and the retinoic acid-related orphan receptors (Ror), including *roraa*, *rorab* (Rorα paralogs), *rorb* (Rorβ), and *rorc*, *rorca*, *rorcb* (Rorγ paralogs). These Ror genes act as transcriptional activators of *arntl* and *clock*/*npas2*, often competing with Nr1d repressors to maintain circadian balance (49, 50).

### Patterns of biological clock gene loss in Perciformes

To assess the status of biological clock genes across Perciformes, we performed a series of pairwise whole-genome alignments and applied the software TOGA (Tool to infer Orthologs from Genome Alignments (51)) to identify orthologs, detect truncating variants, and annotate syntenic gene regions. Genes were considered lost if there are multiple inactivating mutations (e.g., frameshifts, premature termination codons, exon losses, splice site mutations) within the middle 80% of the gene. Genes in loci with fragmented assemblies, lacking syntenic gene regions, with single mutations within the middle 80% of the gene, or with mutations that leave most of the open reading frame intact were given an uncertain status. Aligning genomes to a reference can introduce reference bias due to incomplete gene annotations, assembly artifacts, or species-specific changes to exon boundaries (51). To control for such bias, we aligned all genomes to two separate reference assemblies. In total, 37 genomes assemblies were aligned to reference assemblies of the pikeperch (*Sander lucioperca*, Percidae, Perciformes) and the gilthead seabream (*Sparus aurata*, Sparidae, Spariformes). To capture a representative view of circadian genes across the order Perciformes, we included representative genomes from six of the seven recognized suborders. See **Table S2** for assembly information.

Of the 34 candidate circadian genes, 27 genes were annotated in both reference genomes (*S. lucioperca*, *S. aurata*). Of the seven genes not annotated in these references, one gene, *rorab* (*S. lucioperca*: ENSSLUG00000016495; *S. aurata*: ENSSAUG00010006455), was identified as a previously unannotated gene using a *de novo* annotation approach with Exonerate (52). For the remaining six genes (*arntl1b*, *cry3b*, *cry4*, *per1a*, *csnk1da*, and *nr1d1*), no orthologs were detected with either TOGA or Exonerate in either reference genome and they are presumed to be absent from the genomes of all perciform species included in the study. The absence of these genes is consistent with observed patterns for other members of Acanthomorpha (33).

We detected an unexpectedly widespread pattern of circadian gene loss across Perciformes (**Fig. 1**). These losses were concentrated in two suborders, Notothenioidei and Cottioidei, while other perciform fishes generally retained intact clock genes or have uncertain gene statuses. Within Notothenioidei, the number of gene losses ranged from one in the Patagonian blennie (*nr1d2b*; *E. maclovinus*) and thornfish (*cry1b*; *Bovichtus diacanthus*) to five in several species of Antarctic notothenioids. The most consistent losses in Notothenioidei occurred in *cry1b*, *cry2*, *per2a*, and *per3*, representing shared losses across the Antarctic clade. Although we initially hypothesized that gene losses would be concentrated within Antarctic lineages due to polar seasonality in light/dark cycles, circadian gene loss also occurred outside this group. Notable examples included the channel bull blenny (*Cottoperca trigloides*) with four losses (*cry1a*, *cry3a*, *per2a*, *per3*) and the single losses in *E. maclovinus* and *B diacanthus* described above. Notably, several of these genes lost in sub-Antarctic clades were also independently lost in the cryonotothens, including *cry1b*, *per2a*, *per3,* and *nr1d2b*, which show unique mutational patterns across the phylogeny (**Figs 2**, **S1-S3**).

**Fig. 2.**
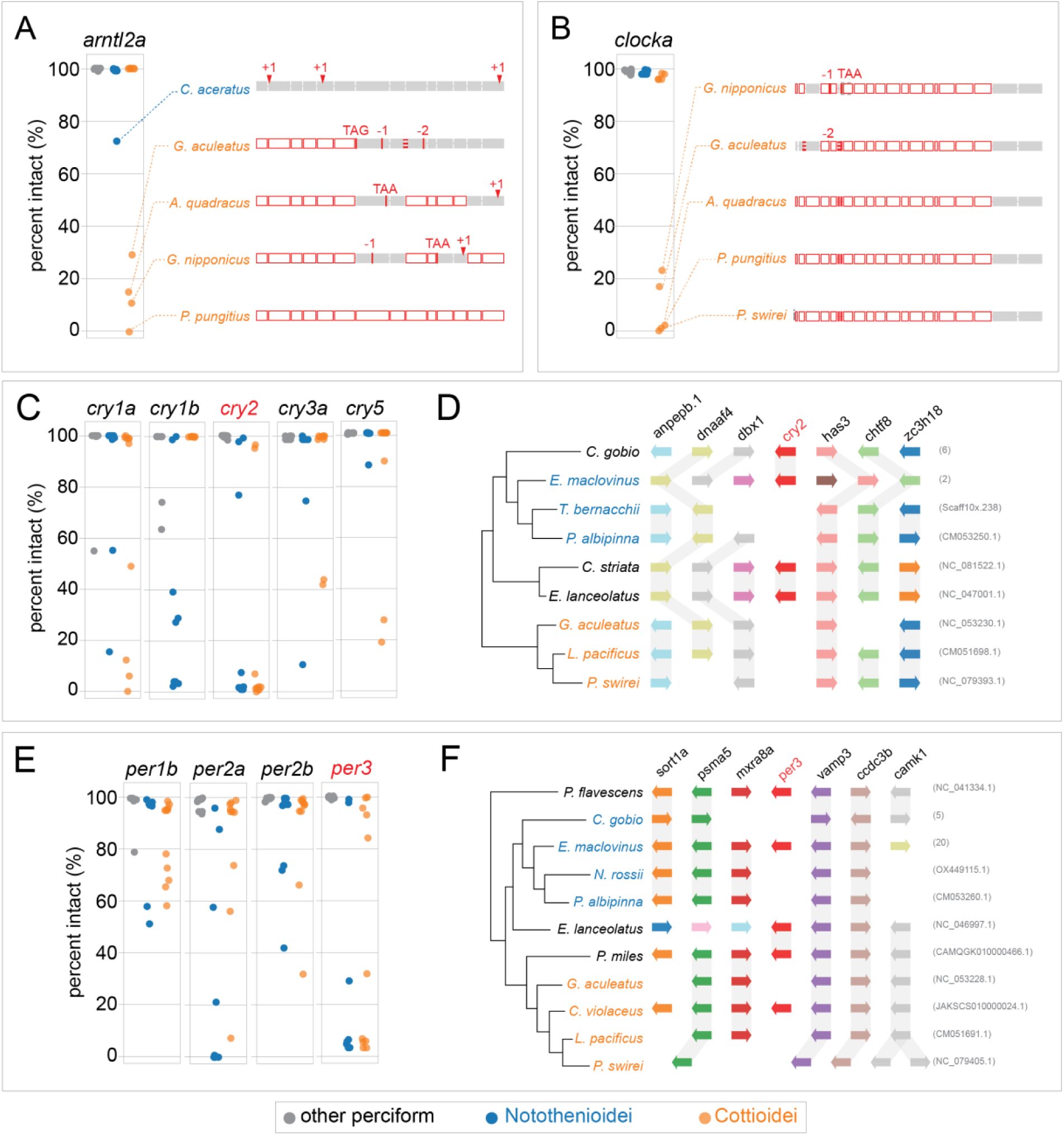
Example mutational variants across central biological clock feedback loop. A–B) Loss patterns of *arntl2* and *clocka*. Plots show the maximum percentage of intact coding sequence (CDS) across all transcript isoforms from pairwise genome alignments to *S. lucioperca* and *S. aurata* reference genomes. Species are grouped as Notothenioidei (blue), Cottioidei (orange), or other Perciformes (gray). Representative mutations across each exon for select species are shown to the right. Exons are shaded gray; deleted or missing exons are white with a red outline. Truncating variants are labeled by their type (e.g., frameshifts as +1 or –1), and splice site mutations are marked with red dashed lines at exon boundaries. C–D) Loss patterns of *cryptochrome* genes. Panel D includes a riparian plot showing conservation of the *cry2* syntenic region across Perciformes. E–F) Loss patterns of *period* genes, with a riparian plot of the *per3* locus in panel F. See **Figs. S1-3** for further mutation information.

In Cottioidei, we also observed numerous gene losses, ranging from no losses in the basally branching sablefish (*Anoplopoma fimbria*) to seven losses (*clocka*, *cry1a*, *cry2*, *cry3a*, *cry5*, *per3*, *rorab*) in the hadal Mariana snailfish (*Pseudoliparis swirei*) (**Fig. 1**). Within the stickleback family (Gasterosteidae), multiple species share losses of *arnl2a*, *clocka*, *cry2*, and *rorab*. Additional lineage-specific losses within Gasterosteidae included *cry1a* and *per2a* in *Apeltes quadracus*, and a shared loss of *per3* in both *Gasterosteus aculeatus* and *G. nipponicus*. Several genes appear to have been lost independently across different Cottioidei lineages, including *cry3a* and *cry5* in both eelpouts and snailfishes; *per2a* in sticklebacks and eelpouts; *rorab* and *clocka* in sticklebacks and snailfishes; and *per3* and *cry1a* in sticklebacks and members of the infraorder Cottales.

There were several key distinctions and similarities between loss patterns in Notothenioidei and Cottioidei. In Notothenioidei, gene losses were concentrated along the Cry/Per axis of the feedback loop, whereas species in Cottioidei have losses in the Clock/Arntl complex in addition to Cry/Per (**Fig. 1**). However, several Antarctic notothenioids also showed uncertain gene status for *clocka*. Four genes (*cry2*, *cry3a*, *per2a*, and *per3*) were lost in both Notothenioidei and Cottioidei. All other gene losses appear to be taxon specific. For example, *cry1b* was lost in individual species of Notothenioidei but not in Cottioidei, while *cry5* was lost in some species of Cottioidei but was not lost within Notothenioidei.

Twelve genes show no clear losses in any perciform species, including *arnt1a*, *clockb*, *npas2*, *per2b*, *timeless*, *csnk1db*, *csnk1e*, *nr1d2a*, *roraa*, *rorb*, *rorca*, and *rorcb*. However, several of these genes have uncertain status in some lineages, which may indicate potential loss of function. Additionally, while many species have lost one of two *cry1* paralogs, there was not conclusive evidence of any individual species losing both *cry1a* and *cry1b*. For example, cryonotothenioids have lost *cry1a* but retain *cry1b.* Conversely, several species have independently lost *cry1b* but not *cry1a*, including *C. trigloides*, *A. quadracus*, *P. swirei*, and *Clinocottus analis*.

In contrast to Perciformes, the five non-perciform species included in this analysis showed virtually no circadian gene losses, despite their greater evolutionary distance from the reference genomes, which could theoretically reduce alignment and annotation accuracy. The only exception was the Atlantic halibut (*Hippoglossus hippoglossus*), which showed a loss in *cry3a*. Notably, the range of *H. hippoglossus* extends into the Arctic, and this species inhabits the highest latitudes among the outgroup species.

### Mutational profiles in clock genes

We found a range of mutations among genes classified as lost, including specific truncating variants and complete deletions (**Figs. 2**, **S1-3**). In many cases, the mutation patterns suggested shared ancestral loss. For example, the first six exons of *arntl2a* were deleted in all examined sticklebacks (**Fig. 2A**). Other cases show strikingly similar patterns that likely reflect convergent evolution based on the species tree topology (18), such as the loss of the first 19 exons of *clocka* in multiple stickleback species and in the hadal snailfish *P. swirei*. Most genes, however, displayed patterns consistent with independent gene loss or drift (**Figs. S1-3**). The most complete gene losses were observed in *cry2* and *per3*, where the majority of species lacked any intact exon sequences (**Fig. 2C-F**). Synteny analysis at the *cry2* (**Fig. 2D**) and *per3* (**Fig. 2F**) loci revealed contagious genome assembly and gene content conservation, supporting loss of these genes. Detailed evidence of loss for each gene in the dataset is provided in **Figs. S1-3**.

### Relaxed selection across clock genes in Perciformes

Given that many species have lost core circadian genes, we hypothesized that other components of the pathway might exhibit more subtle signals of relaxed selection in Perciformes or might be under intensified purifying selection to remove deleterious mutations in the context of a less redundant, streamlined clock gene set. To reduce the impact of reference bias, we only considered a gene to be under relaxed or intensified selection (RELAX (53)) if consistent results were obtained with the gene annotations derived from genome alignments to both reference genomes.

Using an outlier-masked multiple gene sequence alignment and treating both Notothenioidei and Cottioidei as foreground clades, we detected a signal of relaxed selection in nine clock genes (*arntl2a*, *clocka*, *cry1a*, *cry5*, *per1b*, *per2a*, *rorb*, *rorc*, *rorca*; **Fig. 3**; **Table S3**). Except for *per2a*, most of these genes showed only sporadic losses across the phylogeny (**Fig. 1**), and two genes (*rorca* and *rorb*) had no losses at all.

**Fig. 3.**
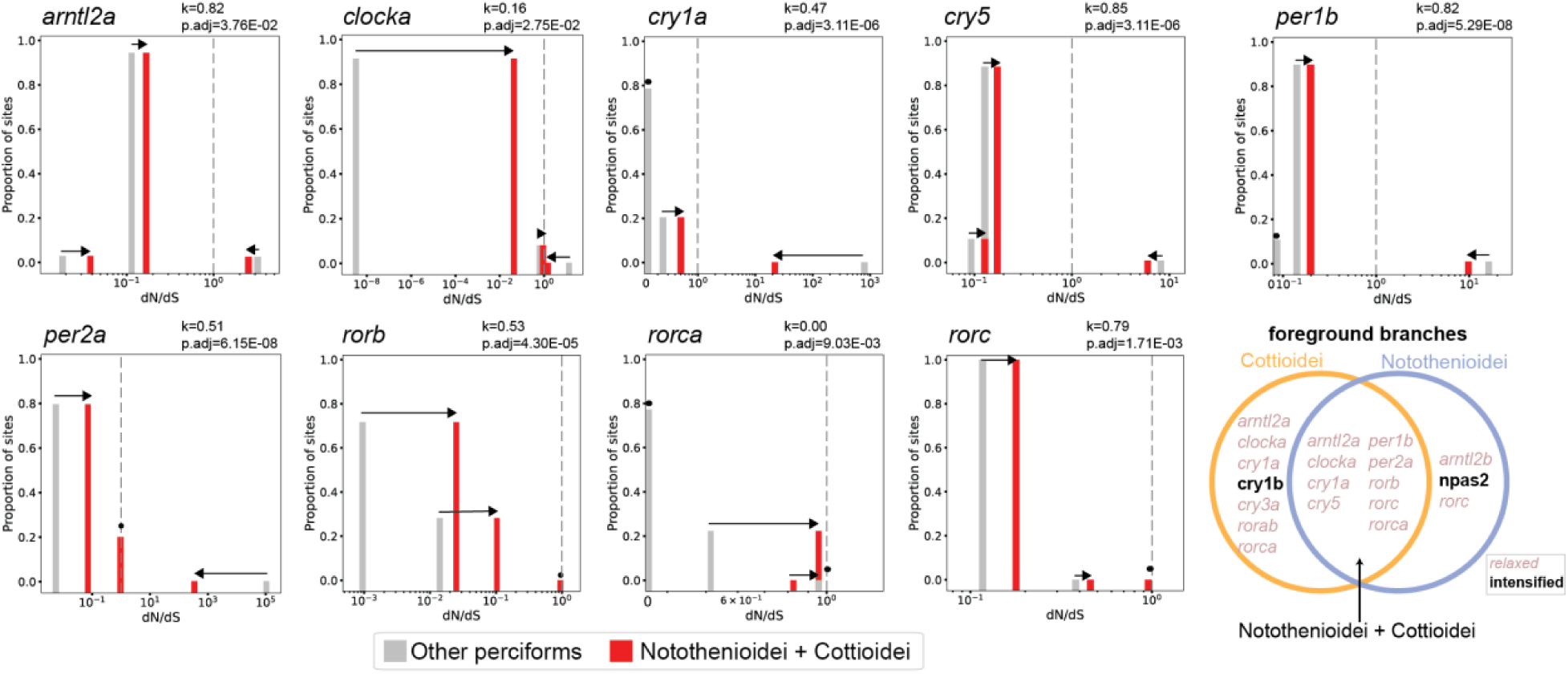
Relaxed selection of biological clock genes in Perciformes. RELAX plots showing changes in selection intensity between foreground branches (both Notothenioidei and Cottioidei) and background branches (other Perciformes). The three ω rate classes are shown: ω₀ (purifying selection), ω₁ (neutral selection), and ω₃ (diversifying selection). Shifts in ω values toward 1 indicate relaxed selection, while shifts away from 1 indicate intensified selection. Plots include only genes with statistically significant selection intensity parameters (k) that were consistent across annotations generated from genome alignments to both *S. lucioperca* and *S. aurata* reference assemblies. K and FDR-adjusted p-value shown relative to *S. lucioperca*. Venn diagram highlights genes under relaxed (pink italics) and intensified (bold) selection when either Cottioidei, Notothenioidei, or both Cottioidei and Notothenioidei are specified as foreground branches. Note, gene entries are listed in multiple locations within the Venn diagram if they were significant in multiple foreground clade selections.

We further investigated whether the observed signatures of relaxed selection were driven by one or both foreground clades. When Notothenioidei was used as the foreground, only three genes showed changes in selection intensity (*arntl2b*, *npas2*, *rorc*; **Fig. 3**, **Table S3**). These genes are intact in most notothenioid species, except for *arntl2b*, which was lost in *Harpagifer antarcticus* (**Fig. 1**, **Fig S3A**). Interestingly, *npas2*, which is retained across all perciform species, was found to be under intensified purifying selection in Notothenioidei.

Cottioidei exhibited changes in selection intensity for seven genes: *clocka*, *cry1a*, *cry1b*, *cry3a*, *rorab*, *rorca*, and *arntl2a*. Some of these genes, such as *clocka* and *arntl2a*, show frequent loss within the suborder. Others, like *cry1a* and *cry3a*, are lost more sporadically, while *cry1b* and *rorca* show no losses at all (**Fig. 1**). Interestingly, *cry1b* is the only cryptochrome gene under intensified purifying selection, and also the only one in Cottioidei without inactivating mutations.

Notably, four genes were only significant with both Notothenioidei and Cottioidei as foregrounds, including *cry5*, *per1b*, *per2a* and *rorb*. In other genes, the statistical significance is likely driven by either Notothenioidei (*rorc*) or Cottioidei (*arntl2a*, *clocka*, *cry1a*, *rorca*) (**Fig. 3**; **Table S3**).

### Circadian gene loss is correlated with latitude

Given the apparent clustering of gene losses and relaxed selection within the Notothenioidei and Cottioidei suborder (**Fig. 4**), we asked whether there is an association between biological clock gene loss and specific environmental variables. Both Notothenioidei and Cottioidei are abundant in polar and high-latitude regions, as well as across a wide depth gradient (18, 20). We hypothesized that these two environments, high latitude and depth, could be associated with relaxed selection in biological clock genes gene loss due to the seasonally arrhythmic or absent light/dark cycles that characterize these environments respectively (**Fig. 5A, B**). Because our dependent variable (count of circadian gene losses) is discrete and prone to zero inflation, we tested multiple models and distributions with a combination of counts, log-transformed counts, and binary losses to find the best fit to our data resulting in five mixed models (MCMCglmm) and one general linearized model (Phyloglm). All six models investigated show a significant positive relationship between mean latitude and circadian gene loss with the Gaussian distribution run on log-transformed losses having the best overall fit (**Fig. 5**). Only one of the models, the Gaussian distribution applied to loss count data, found a significant relationship between depth and circadian gene loss, with all other models having a p-value > 0.05.

**Fig. 4.**
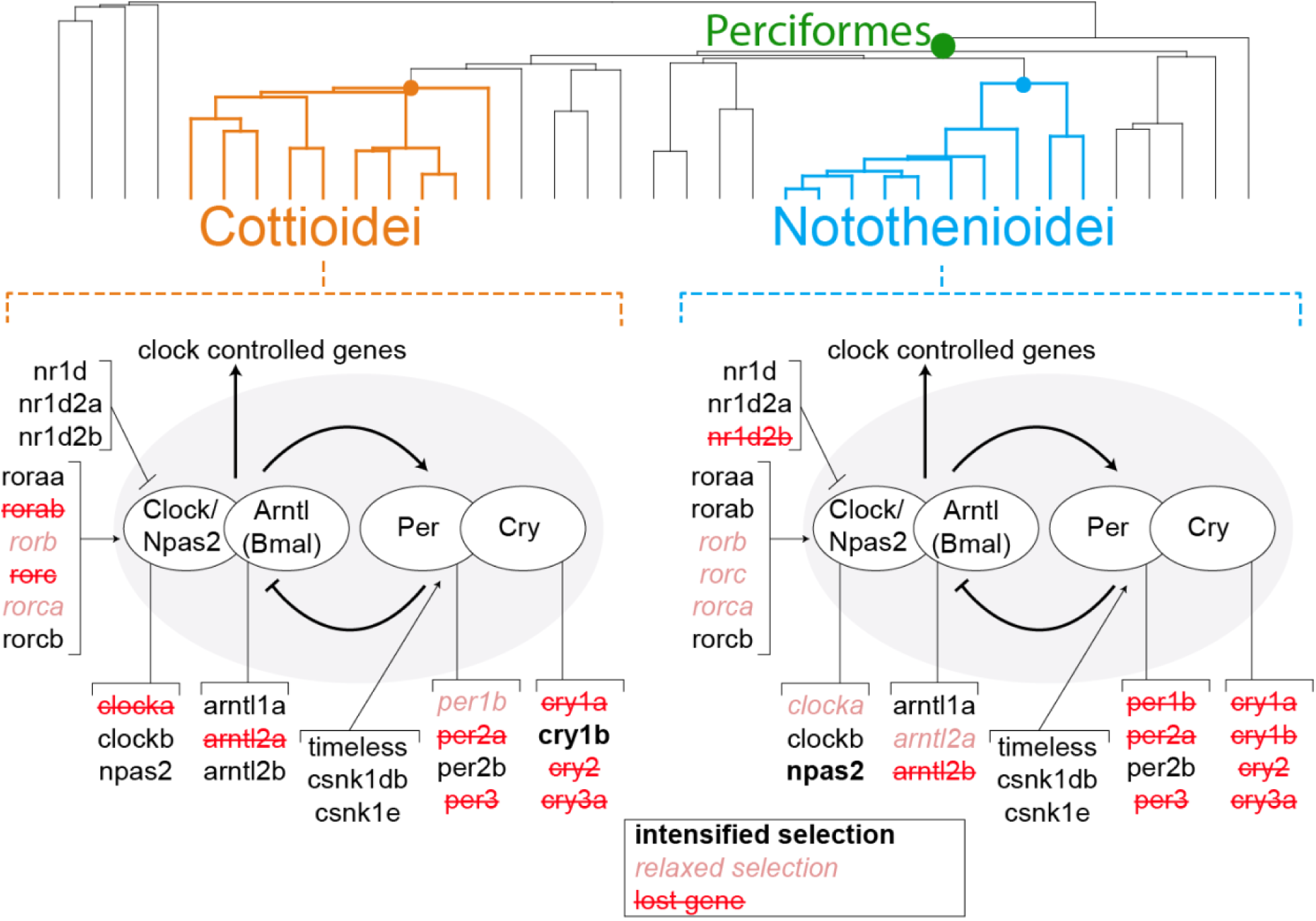
Summary of biological clock gene status across Notothenioidei and Cottioidei. Red strikethrough text indicates gene loss in at least one species from each group. Note individual fishes within each clade may possess intact versions of these genes (Fig. 1). Pink italicized text indicates relaxed selection within the group, while black bold text indicates intensified selection.

**Fig. 5.**
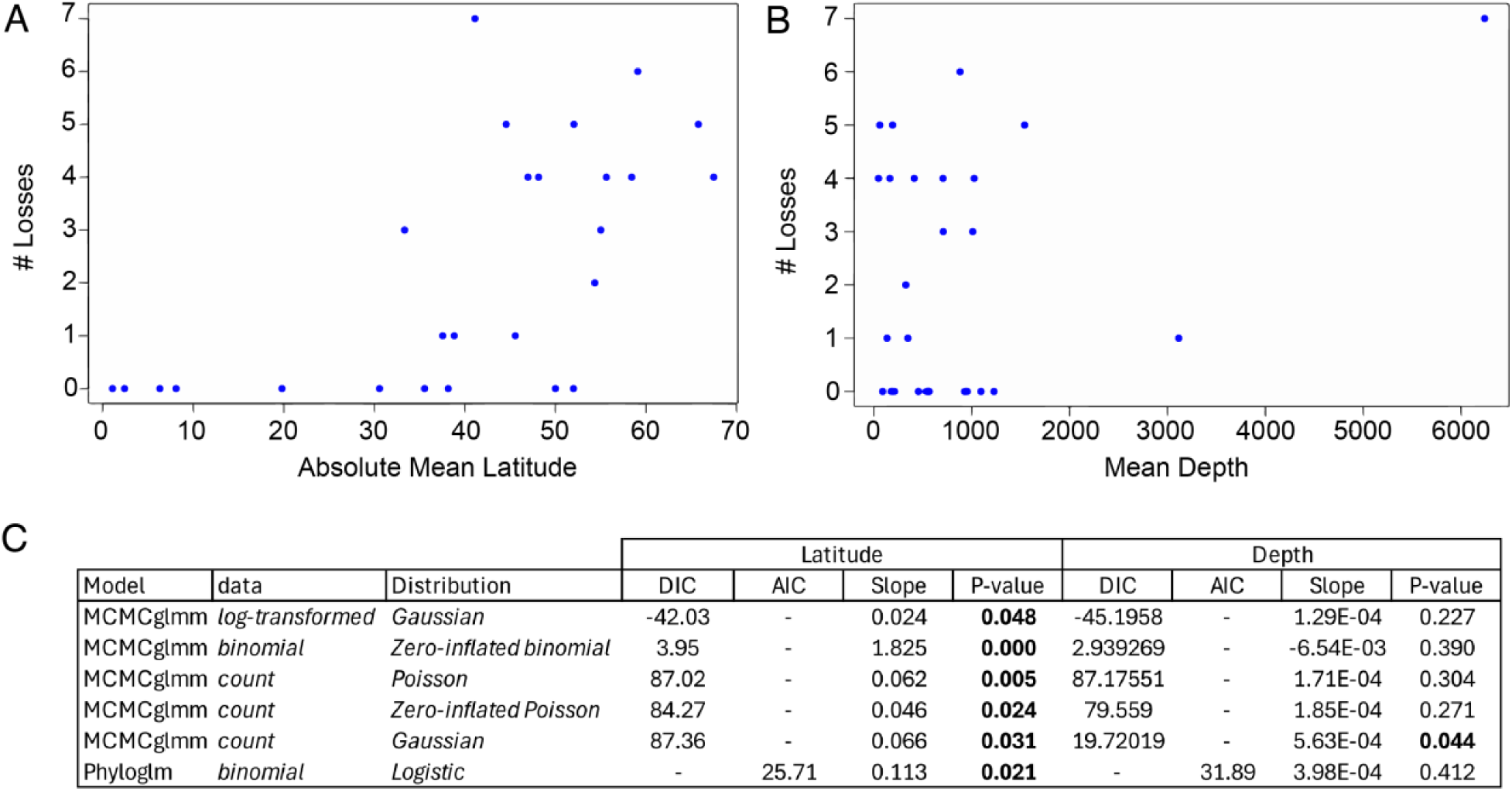
Biological clock gene loss correlates with latitude. Scatter plots show (A) the number of circadian gene losses plotted against the absolute value of mean species latitude, and (B) gene losses plotted against mean species depth. Species distribution data are from AquaMaps (88). C) Results from multiple phylogenetic models testing the association between gene loss and latitude. Statistically significant results are shown in bold.

### Apparent absence of circadian rhythms in the metabolic rate of Notothenioidei

The observations of circadian rhythms in Perciformes are sparse, but have identified seasonal and inter-individual variability in diel activity patterns (30–32). Given the widespread patterns of gene loss and relaxed selection in the biological clock, we asked whether these fishes had evidence for circadian rhythms in metabolic rate and whether this was restricted to polar clades or if it was shared with temperate species. Leveraging the quiescent traces of standard metabolic rate (e.g. minimum maintenance metabolism), we analyzed continuous patterns of metabolic rate over a 48-hour window from three species of Notothenioidei. Over two diurnal cycles, European seabass (*Dicentrarchus labrax*), a temperate acanthuriform species, exhibited a metabolic oscillation at an interval of 25 ± 0.83 hours (**Fig. 6A**). Intriguingly, Notothenioidei, including sub-Antarctic (*E. maclovinus*) and two Antarctic species (*Pseudochaenichthys georgianus* and *C. aceratus*), all had no oscillation in their metabolic profiles over the two diurnal cycles (**Fig. 6B-D**). Instead, the fishes of Notothenioidei had a consistently quiescent metabolic profile situated at a steady baseline level.

**Fig. 6.**
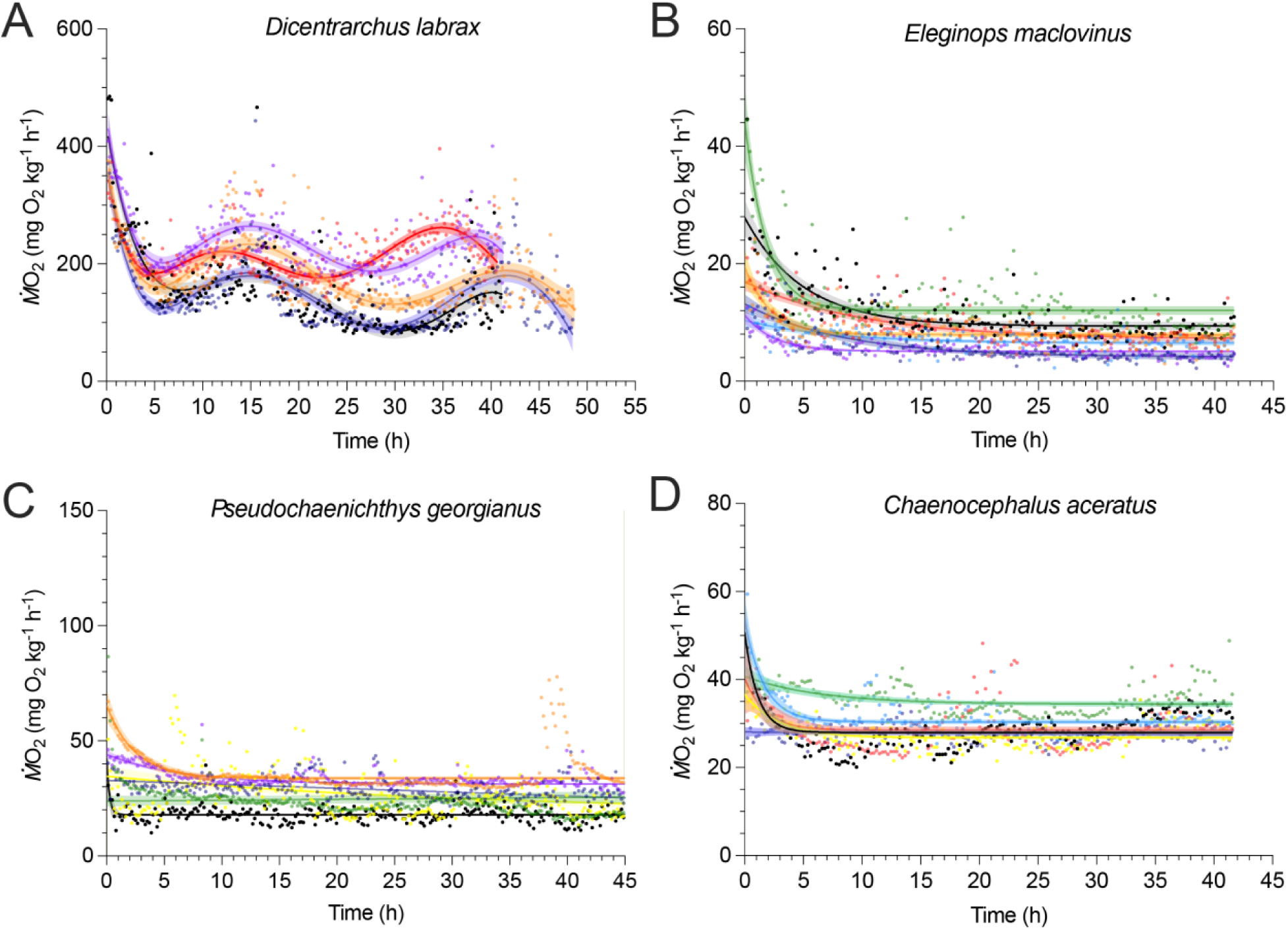
Absence of circadian metabolic profiles in Notothenioidei. Metabolic profiles across two diurnal cycles in four representative species of Percomorpha. *D. labrax* (A) is a temperate acanthuriform species. *E. maclovinus* (B), *P. georgianus* (C), and *C. aceratus* (D) are notothenioid species within the Notothenioidei clade. Colors represent different individuals. Dots indicate measured aerobic metabolic rates. Solid lines show the fitted regression models, with shaded areas representing 95% confidence intervals. A sixth-order polynomial model was used for *D. labrax*, and a one-phase decay model was applied to *E. maclovinus*, *P. georgianus*, and *C. aceratus*.

## Discussion

Our findings indicate that loss and relaxed selection in biological clock genes occur along a latitudinal gradient and has evolved convergently both within and between two perciform suborders, Notothenioidei and Cottioidei. Though limited, we find no evidence of circadian rhythm of metabolic rate in Notothenioidei, indicating that these gene losses correspond to likely phenotypic variation. Plasticity in the circadian patterns of activity may constitute an evolutionary predisposition that facilitates survival in arrhythmic environments.

### Streamlining the perciform biological clock gene set

The gene loss patterns we identified in Notothenioidei and Cottioidei align with previous reports of biological clock gene disruptions in notothenioids (25, 26, 54), deep-sea snailfishes and eelpouts (28, 55), as well as the loss of *per3*, *arntl2a*, and *clocka* in the three-spined stickleback (33, 39). We expand on these findings by showing a broader, latitude-correlated pattern of circadian gene loss across Perciformes (**Figs. 1**, **4**, **5**).

Our data indicate distinct preservation patterns in core biological clock gene paralogs within Perciformes, potentially highlighting reliance on specific clock genes over others. Many infrequently lost genes have pleiotropic functions outside the clock. For example, Cry5, Cry-dash, and Timeless contribute to DNA repair, and retinoid-related orphan receptors (Ror) are involved in immune, metabolic, and developmental processes (43, 47, 56). These broader functions likely explain their retention in Perciformes. The consistent presence of *clockb*, *npas2*, and *arntl1a* also suggests essential roles. In contrast, the species-specific retention of either *cry1a* or *cry1b* suggests some functional redundancy between these paralogs. One of the most consistent losses is *per3*, which is absent in multiple species of Notothenioidei and Cottioidei. This gene is frequently lost across teleosts (33), including in Atlantic cod (*Gadus morhua*), northern Pike (*Esox lucius*), three-spined stickleback (39), and in many salmonids (40, 57), suggesting a diminished role in fish circadian regulation. Still, *per3* remains under intensified purifying selection in Atlantic salmon (*S. salar*), and brown trout (*Salmo trutta*) (40, 57). In contrast, *per1b* and at least one copy of *per2* are relatively well conserved across teleosts (40), including in our dataset (**Fig. 1**).

The gene *npas2* is retained across all species in our dataset and is under intensified purifying selection in Notothenioidei (**Figs. 1**, **3**). The Npas2 protein has two PAS (Period–Arnt–Single-minded) domains that bind heme, are sensitive to carbon monoxide, and dimerizes with Arntl (Bmal) in a redox-dependent manner (58, 59). These features may be important for other functions in notothenioids, which potentially face high cellular oxidative stress in the oxygen-rich Southern Ocean and exhibit expansions of oxidative stress-related gene families within the genomes of multiple species (60).

This study focused on gene loss within the core biological clock. However, peripherally important circadian genes and downstream targets are likely under varied selection regimes. Given the repeated invasion of species of Notothenioidei and Cottioidei into arrythmic habitats, this is a promising clade for “forward genomic” and genotype-to-phenotype mapping strategies to identify novel circadian-related genetic components through the detection of patterns of convergent molecular evolution (e.g., (61–63)).

### Lack of metabolic rhythms across Notothenioidei and Cottioidei

The three notothenioid fishes measured here show no evidence of metabolic oscillation. In contrast, *D. labrax*, which retains all core circadian genes tested (**Fig. 1**), exhibits a robust metabolic rhythm that persists once individuals are placed in constant darkness. Notothenioids, however, maintain a low and steady metabolic rate, consistent with findings of sticklebacks held in the dark and Arctic sculpins in constant lighting (**Fig. 6**) (30, 31). The absence of rhythms in our dataset is also consistent with the apparent arrhythmia of *N. coriiceps* held in sea pens (29).

Reports of species entirely lacking circadian rhythms are rare, even in cave environments, and studies often do not exhaustively test all possible zeitgebers or potential rhythmic traits (14). For example, Somalian cavefish (*Phreatichthys andruzzii*) appear to have no overt rhythms, but are nonetheless capable of being entrained on food (64). There is evidence for rhythms in fish clades found well below the penetration of sunlight (1,000m) (e.g., (65, 66)), and many deep-sea invertebrates have circatidal rhythms (67). Within Cottioidei, an intertidal eelpout (*Zoarces vivparius*) maintains circatidal swimming rhythms in captivity before transitioning to photic cues (68).

Given these results from other species, it seems unlikely that notothenioids do not exhibit circadian rhythms of any sort, and that rhythms may vary by season. Additionally, there may be distinctions between benthic and pelagic species. For example, the pelagic Antarctic silverfish (*Pleuragramma antarcticum*) and Antarctic toothfish (*Dissostichus mawsoni*) both show evidence of diel vertical migrations, though the migration of *D. mawsoni* was weaker and more closely followed movements of their prey (*P. antarcticum*) than light intensity (69). Notably, reports on the extent or existence of diel migrations in pelagic notothenioids seem to vary by method and location (70–72). Further research is needed to assess the presence and potential seasonality of circadian rhythms across a diverse spectrum of notothenioids, including multimodal analyses of behavior, hormones (e.g., melatonin), gene expression, and testing of alternative zeitgebers to determine whether these fishes exhibit rhythmicity or are capable of entraining clocks to specific environmental cues.

### Clocks and survival in high latitudes and the deep sea

Fish clades with the fastest speciation rates often include species found in both polar and deep-sea environments, suggesting shared adaptations for thriving in these extreme habitats (20). While most studies focus on cold temperatures or high hydrostatic pressures in these environments, the breakdown of environmental zeitgebers and their impact on engrained circadian mechanisms is also likely a major adaptive barrier (16). Perciformes are overrepresented in both high-latitude and deep-sea ecosystems, including polar oceans and some of the deepest fish-inhabited regions (18, 20, 73, 74) (**Table S4**). In such environments, responding opportunistically rather than anticipating cyclic cues may be advantageous, effectively keeping physiological systems active at a low baseline to exploit unpredictable conditions. We found that circadian gene loss increases with latitude, suggesting relaxed selection for maintaining a 24-hour clock at higher latitudes (**Fig. 5**). In contrast, we found no clear correlation between gene loss and depth (**Fig. 5**), but with more deep-sea genomes this may change. Further, many deep-sea species are migratory, span wide depth ranges, or lack precise depth data due to limitations in catch records, complicating analyses based on depth.

The sub-Antarctic notothenioid *E. maclovinus* had comparatively few circadian gene losses compared to Antarctic notothenioids but shows the same lack of metabolic oscillation as *P. georgianus* and *C. aceratus* (**Figs. 1**, **6**). This suggests that reduced reliance on circadian rhythms may be a broader trait of Notothenioidei and could have supported their adaptive radiation. As with circadian rhythms, low skeletal density is another trait shared between *Eleginops* and the Antarctic clade (75). Reduction in skeletal density is a key buoyancy adaptation in notothenioids, which lack swim bladders, and has been proposed as a historical contingency that could have enabled the clade to more rapidly diversify into midwater niches (76). Since disruption of circadian rhythms is considered a major barrier to polar adaptation (16), diminished circadian dependence may be an overlooked trait that facilitated this evolutionary expansion. Non-polar perciforms show signs of diel flexibility. In addition to the stickleback example mentioned above (30), the slimy sculpin (Cottus cognatus; Cottioidei) in Lake Ontario (late 43°N) shows evidence of diel rhythms in feeding as juveniles at higher water depths (35 m) but has arrhythmic feeding patterns as adults at 75 m (77). Given this flexibility, perciform clades may be primed to thrive in these arrhythmic environments.

### Summary

We find that many core biological clock genes are disrupted and under relaxed selection in two perciform clades common to high-latitude and deep-sea environments: Notothenioidei and Cottioidei. The extent of gene loss correlates with latitude, with polar species showing more loss than temperate lineages. We also show that both Antarctic and sub-Antarctic notothenioids lack daily metabolic rhythms, instead exhibiting steady-state baseline metabolism like that observed in Cottioidei under constant lighting. While high-latitude and deep-sea habitats impose diverse selective pressures, our findings support the idea that altered circadian regulation is a key adaptation to these environments. Because sustained circadian disruption can be harmful and lead to disease in vertebrates, understanding how these species maintain circadian plasticity and healthy physiological function may offer insight into resilience to circadian-related pathologies

## Materials & Methods

### Genome alignment, ortholog calling, and determination of gene status

To assess the presence or absence of circadian genes, we employed a whole genome alignment approach that leverages both gene sequence content and syntenic gene context to identify and characterize orthologs across Perciformes. We pairwise aligned a total of 32 perciform and five outgroup genome assemblies to two different reference assemblies, *Sander lucioperca* (GenBank ID: GCA_008315115.1) and *Sparus aurata* (GCA_900880675) (**Table S2**), using the program *make_lastz_chains* (v 1.0.0) (https://github.com/hillerlab/make_lastz_chains) (78–80). These aligned genomes included 10 species in the suborder Notothenioidei (*Cottoperca trigloides*, *Bovichtus diacanthus*, *Eleginops maclovinus*, *Trematomus bernacchii*, *Notothenia rossii*, *Harpagifer antarcticus*, *Pogonophryne albipinna*, *Gymnodraco acuticeps*, *Pseudochaenichthys georgianus*, *Chaenocephalus aceratus*), 10 species in the suborder Cottioidei (*Gasterosteus aculeatus*, *Gasterosteus nipponicus*, *Apeltes quadracus*, *Pungitius pungitius*, *Cebidichthys violaceus*, *Lycodopsis pacificus*, *Cyclopterus lumpus*, *Pseudoliparis swirei*, *Clinocottus analis*, *Anoplopoma fimbria*), four species of the suborder Percoidei (*Echiichthys vipera*, *Gymnocephalus cernua*, *Perca flavescens*, *Etheostoma perlongum*), four species of the suborder Serranoidei (*Centropristis striata*, *Hypoplectrus puella*, *Epinephelus lanceolatus*, *Epinephelus cyanopodus*), three species of the suborder Scorpaenoidei (*Synanceia verrucosa*, *Pterois miles*, *Sebastes schlegelii*), one species of the suborder Triglioidei (*Chelidonichthys spinosis*), and five non-perciform outgroups (*Dicentrarchus labrax, Thunnus maccoyii, Hippoglossus hippoglossus, Oreochromis aureus, Melanotaenia boesemani*).

To identify orthologs, annotate genes, and determine the gene status for the query genomes, we ran TOGA (v. 1.1.0-blue) (Tool to infer Orthologs from Genome Alignments (51)) on all pairwise alignment chains. TOGA identifies orthologs between the reference and query genome and analyzes the middle 80% of the query’s open reading frame to determine the gene status. Each gene was classified as intact (I), partially intact (PI), lost (L), or uncertain loss (UL). Briefly, a gene was considered lost if it contained multiple inactivating mutations (e.g., frameshifts, premature stop codons, exon deletions, or splice site disruptions) within the central 80% of the protein-coding sequence (CDS) across all transcript isoforms. Genes with only a single inactivating mutation in this region were categorized as uncertain losses (UL), as such cases may result from limited exon conservation, gene annotation errors, or assembly artifacts (81). Genes with inactivating mutations outside of the middle 80% are considered partially intact (PI).

Gene annotations based on pairwise genome alignments are prone to reference bias (51). To address this, we compared the status of each gene in each species relative to both reference assemblies. We used the TOGA orthology inferences generated from a pairwise alignment between the *S. lucioperca* and *S. aurata* genome assemblies to map and compare gene statuses across the different multiple genome alignment datasets. For any discrepancy we conservatively selected the more intact status between the two reference alignments.

It is also possible genes are lost within the reference species. To determine the status of circadian genes in the reference species we ran the assembled genomes through the pipeline described above, but with the Nile tilapia (*Oreochromis niloticus,* Cichliformes; GCA_013358895.1) as the reference genome. Notably, aligning to multiple reference assemblies improved the proportion of orthologous genes labeled as intact (I/PI) across the entire dataset, from an average of 67.6% in *S. lucioperca* and 70.9% in *S. aurata* alignments respectively to 73.7% intact (I/PI) in the merged dataset (**Table S5**).

Genome-wide, we identified orthologs, regardless of intact status, for ≥ 90% of all reference-annotated genes in all species examined, except for the thornfish *Bovictus diacanthus* (Notothenioidei), which showed slightly lower recovery rates (87% for *S. lucioperca*, 89% for *S. aurata*). Several genes in our candidate list were not explicitly annotated within the *S. aurata* or *S. lucioperca* genome assembly. To determine if these genes were in fact present but unannotated within these species, we used data from a distant relative, the zebrafish (*Danio rerio*, Cypriniformes; GCA_000002035.4). CDS and protein sequence for each candidate gene were extracted from *D. rerio*. We then used the gene model mapper GeMoMa (v 1.9) (82) to infer the gene annotation in our query species and produce an annotation file in gff format with regions that are a likely match to the reference gene. We then used Exonerate (v 2.2.0) (52) to validate the match between regions annotated by GeMoMa and the protein sequence of the query circadian genes. Any query circadian genes that were not identified in the analysis were considered missing and excluded from the study.

### Relaxed selection

To complement our dataset of lost genes we sought to identify circadian genes that are under relaxed selection, but do not meet the status of “lost” as implemented by TOGA. To select representative transcript isoforms for selection analyses within each species, we used ISOSEL (v 1.0) (83), which uses a phylogenetic analysis to score isoforms based on overall conservation across the dataset. A multiple codon alignment was then generated using MACSE (v 2.07) (84, 85). To control for potential artifacts, such as alignment errors or inclusion of non-exonic sequence due to reference bias during annotation, we identified and masked outlier sequences using TAPER v 1.0.0 (86). Unlike block-based alignment trimming tools, TAPER detects and masks stretches of unusually divergent sequence within individual species based both on the degree of divergence at specific alignment sites and in the context of the rest of the sequence. We also used BUSTED-E (v 4.5) in HYPHY (v 2.5.63) to identify and mask specific codons with exceptionally high ω values (>100), which are indicative of alignment issues (87). Phylogenetic patterns of relaxed or intensified selection were then inferred using RELAX (v 4.1)(53). The tree topology used for this analysis was pruned from Rabosky et al (18). Foreground lineages included both tips and ancestral branches with respect to Notothenioidei and/or Cottioidei. We ran this pipeline on circadian genes that were annotated based on alignment to both *S. aurata* and *S. lucioperca*, as both the gene annotations reconstructed from reference genomes and the selected representative gene isoforms for each species may contain distinct sequences across the independent RELAX runs.

### Correlation of circadian gene loss and the environment

To test if there is a statistical relationship between circadian gene loss and latitude or depth, we fit multiple models to our dataset. We used a count of circadian genes with a status of “lost” as our dependent variable and latitude or depth as our explanatory variable. We sourced latitude and depth data from the website Aquamaps (aquamaps.org), which aggregates species collection data from multiple biodiversity databases (88). We calculated a mean latitude and depth from all observations for the 26 perciform species for which data was available (**Table S4**). We used the absolute value of latitude in our analysis to account for species in both hemispheres. We fit multiple models to our data to account for a discrete dependent variable and the possibility of zero-inflation. We first fit a general linearized model to the data using the R package phyloglm (v 2.6.5) and used a variance-covariance matrix derived from our phylogeny to account for evolutionary history (89–91). We converted loss counts to binary (zero for no lost genes, one for one or more lost genes) and fit a Poisson distribution. To account for the possibility of zero-inflation and to test additional distributions while still accounting for the evolutionary relationship, we also used a Bayesian phylogenetic generalized linear mixed models in the R package MCMCglmm (v 2.36) (92). We ran the MCMC for 500,000 iterations with a 10% burn in and a thinning interval of 40. For all models, we used weakly informative priors (V = 1, nu = 0.002) to minimize their influence on posterior probabilities. To control for the presence of zeros in our data, we tested models with standard counts, log-transformed counts, and binary losses. We used zero-inflated Poisson (ZIP) and standard Poisson distributions with loss counts, a Gaussian distribution with log-transformed losses and a count of losses, and zero-inflated binomial (ZIB) for binary losses.

### Collection and housing of experimental animals

Notothenioid species were collected from regions of Low Island (63° 25′ S; 62° 10′ W) and Dallmann Bay (64° 10′ S; 62° 35′ W). The fish gears are benthic otter trawls deployed from US *ARSV Laurence M. Gould* during the austral fall and winter of 2023. Fish were held in circulating seawater tanks onboard the ship. Fish were then transferred to the aquarium at the Palmer Station (US Antarctic Research Program). At the station, the fish were held in tanks with circulating seawater at 0.1 ± 0.5◦ C. All tanks were equipped with oxygen diffusers and blocks of frozen seawater were added as needed to maintain the temperature. All experimental procedures were approved by the University of Alaska Institutional Animal Care and Use Committee (IACUC; protocols 247598-11 and 570217-9). In the austral fall of 2024, juvenile *Eleginops maclovinus* (Cuvier, 1830), were captured in Reloncavi Fjiord (12°C). The fish were held in circulating tanks (8°C) at Los Lagos University for two months before their transportation to Laboratorio Costero de Recursos Acuáticos Calfuco (Universidad Austral de Chile). In the laboratory, the fish were acclimated to 4°C, bringing the thermodynamics efforts on the fish physiology closer to that of the Antarctic notothenioids. The temperature acclimation started with the reduction of the water temperature at a pace of ∼1.75 °C per day. The fish were then held at this temperature for 2.5 weeks. Animal collection, care and use were approved by permit from the Universidad Austral de Chile and UAF IACUC (above).

A cohort of European sea bass (*Dicentrarchus labrax*) was acclimated to laboratory conditions in a 2000-L indoor tank for 2 months, during which they were fed *ad libitum* twice weekly (Le Gouessant, Lambale, France). A sub-cohort of juvenile European sea bass was distributed between two 500-L tanks. These tanks were supplied with open-flow seawater. The dissolved oxygen level was maintained above 90% air saturation (> 8.2 mg L^-1^). After completing the metabolic rate measurement trial, all fish were placed in a recovery aquarium before being returned to the holding tank after one hour. Fish holding and experimental procedures followed the guidelines of animal care rules and regulations in France (Apafis 2018040916374437).

### Aerobic metabolic rate

We measured the circadian rhythm of the metabolic rate using an automatic intermittent-flow aquatic respirometry system. The respirometry system measured four individuals simultaneously, each within their own chamber. Hence, the metabolic profiles can be obtained for every individual. The automation features of the system enabled undisturbed and continuous measurements for 48 hours. To avoid the effects of digestion, spontaneous activities, and movement, the animals were fasted for 48 hours to reach a post-absorptive state. The metabolic rate of the animals was then measured in a dark and quiescent environment. Thus, only the innate rhythm is manifest in the metabolic profiles. The details of the metabolic rate calculation and the metabolic measuring techniques can be found below and in previously published studies using the same type of respirometry system (93–95).

The respirometry system used Loligo®-type (Loligo®Systems, Denmark, loligosystems.com) respirometer chambers. The size of the chamber was matched to the fish size (water volume : fish ratio ∼ 27 : 1) to optimize detection sensitivity for the change in water dissolved oxygen (DO) concentration inside of the respirometer due to fish oxygen uptake when the respirometer was in the closed mode. All fish were fasted for at least two days before being placed in the respirometer, where the rate of oxygen uptake (*Ṁ*O_2_) was continuously monitored for ∼two days, and only the fish that were not visually agitated were included in the final analyses. All four chambers were immersed in a temperature-controlled seawater bath, which was connected via a pump (Eheim 600) to a gas exchange column that delivered aerated water to the respirometers in the open mode. DO was maintained above ∼85 % saturation throughout the protocol.

Background respiration of each empty respirometer chamber was measured for 20 min before and after each trial and found to be negligible relative to the minimum maintenance metabolic rate of notothenioid species (< 1%, where disinfected seawater was used). For European sea bass, the background respiration in 25 °C seawater exceeded this 1% threshold and the fish’s *Ṁ*O_2_ was corrected by subtracting the background *Ṁ*O_2_ value. Background respiration was minimized by thoroughly disinfecting the entire apparatus with sodium hypochlorite (Performance bleach, Clorox in 1000 ppm) for 30 min (European sea bass study).

*Ṁ*O_2_ was continuously and automatically monitored on-line using computer software (AquaResp v.3, Denmark, Aquaresp.com) that processed water DO measurements in the respirometers (a 1 Hz sampling rate) from an optical oxygen probe associated with each respirometer (Robust Oxygen Probe OXROB10, Pryoscience, pyro-science.com). The optodes were calibrated to 0% saturation (water saturated with sodium sulphite and bubbled with nitrogen gas) and 100% saturation (fully aerated water) at the start of each experiment. *Ṁ*O_2_ values were calculated whenever the respirometers were sealed. The measurement cycles (flush period, stabilization period and sealed period) are 65-208-600 for *E. maclovinus*; 80-200-330 for *P. georgianus*; 80-200-600 for *C. aceratus*; and 120-60-420 for *D. labrax*.

The slope of the decrease in DO over time met a minimum requirement for linearity (*i.e.*, R^2^ > 0.9) to calculate *Ṁ*O_2_). The quality of PO_2_ traces was checked as described by Chabot et al (96). Metabolic rate measurements are directly calculated from AquaResp software using the conventional sequential algorithm (**Eqn.1**.)

### Aerobic metabolic rate calculations

Aerobic metabolic rate was calculated with the following equation:

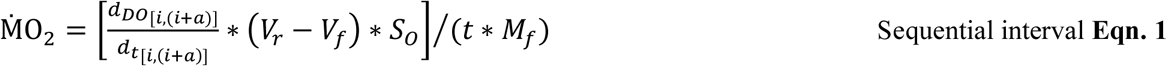

where units for *Ṁ*O2 are mg O2 h^-1^ kg^-1^, 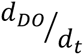 is the change in O_2_ saturation over time, *V_r_* is the respirometer volume, *V_f_* is the assumed fish volume, *S_o_* is the solubility of O_2_ (calculated by AquaResp v.3 software) in the experimental temperature, salinity and atmospheric pressure, *t* is a time constant of 3600 s (per hour), *M_f_*is fish mass, *a* is the sampling window duration (s), *i* is 1 DO sample forward from the end of previous sampling window at a set sampling frequency of 1 Hz.

### Modeling of the metabolic profiles

The modeling of the circadian rhythm of D. labrax using the sixth order polynomial model (**Eqn. 2**):

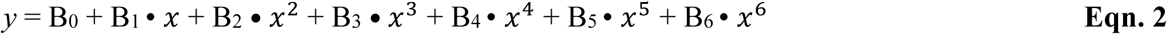

where B_0_, B_1_, B_2_, B_3_, B_4_, B_5_, B_6_ are the best fitted coefficients.

The modeling of the circadian rhythm of *Nototheniodei* use one phase decay model (**Eqn. 3**):

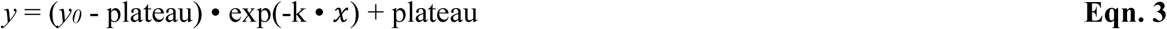

where y_0_ is the intercept, plateau is the y value extrapolated at infinite distance at x-axis, k is the rate constant.

The regression analyses were conducted in Prism v.10 (GraphPad Software, USA, graphpad.com).

## Acknowledgements

This work was supported by the National Science Foundation (NSF) grant OPP-2324998 and by a National Institutes of Health (NIH) grant 1R35GM150590 to J.M.D. This work as also supported by an NSF postdoctoral fellowship OPP-2420167 to D.B.W. Metabolic rate data were collected by Y.Z. in collaboration with Dr. Kristin M. O’Brien (University of Alaska Fairbanks) under NSF PLR-1954241 (K.M.O.). YZ is supported by a Postdoctoral Fellowship of the Natural Sciences and Engineering Research Council of Canada (NSERC PDF - 557785 – 2021) followed by a Banting Postdoctoral Fellowship (202309BPF-510048-BNE-295921) of NSERC & CIHR (Canadian Institutes of Health Research). This Antarctic fieldwork was made possible by the support of the captain and crew of the *ASRV Laurence M Gould*, by the staff at Palmer Station, and by personnel at the Office of Polar Programs at the National Science Foundation (NSF). This work was completed in part with resources provided by the Research Computing Data Core at the University of Houston.

## Author contributions

D.B.W., Y.Z. and J.M.D. designed and performed research; D.B.W., Y.Z., and J.M.D. analyzed data; and D.B.W. and J.M.D. wrote the paper; D.B.W., Y.Z. and J.M.D. provided critical revision of the manuscript.

---

**Fig. S1.**
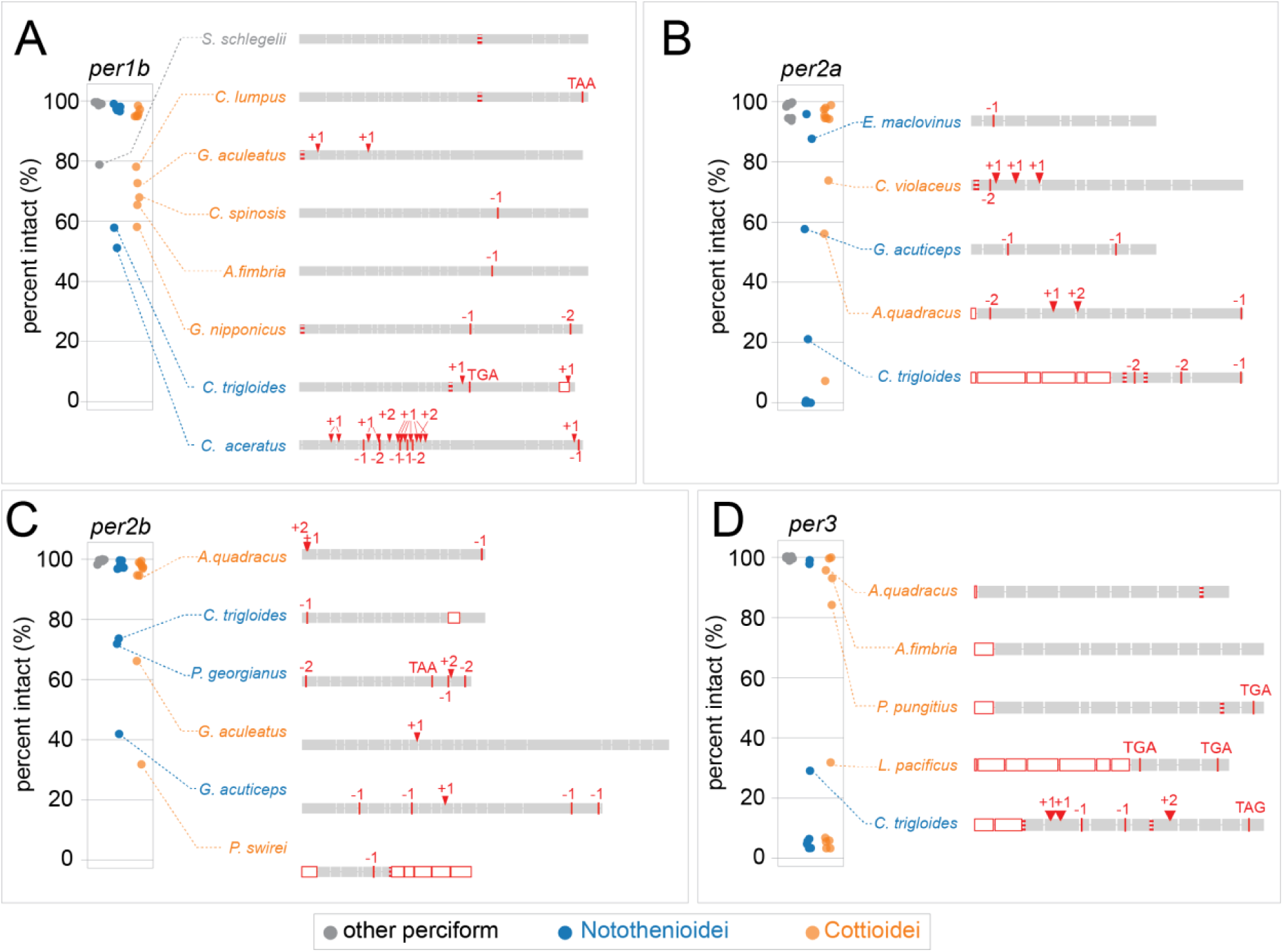
Example mutational variants across *period* genes. A) *per1b*, B) *per2a*, C) *per2b*, D) *per3*. Plots show the maximum percentage of intact coding sequence across all transcript isoforms annotated via pairwise genome alignments to *S. lucioperca* and *S. aurata* reference genomes. Species are grouped as Notothenioidei (blue), Cottioidei (orange), or other Perciformes (gray). Representative mutations across each exon for select species are shown to the right. Exons are shaded gray; deleted or missing exons are white with a red outline. Truncating variants are labeled by their type (e.g., frameshifts as +1 or –1), and splice site mutations are marked with red dashed lines at exon boundaries.

**Fig. S2.**
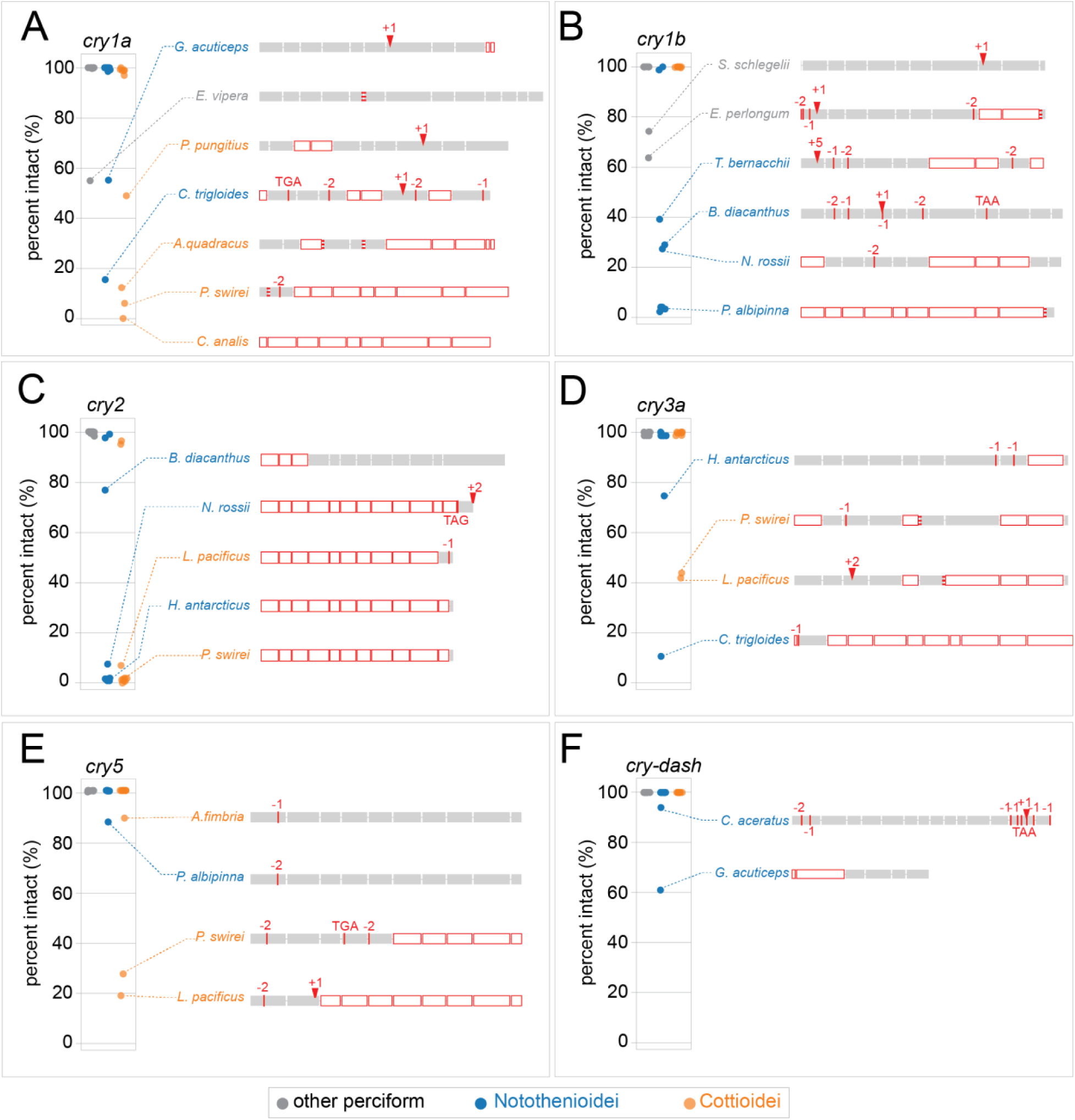
Example mutational variants across *cryptochrome* genes. A) *cry1a*, B) *cry1b*, C) *cry2*, D) *cry3a*, E) *cry5*, F) *cry-dash*. Plots show the maximum percentage of intact coding sequence across all transcript isoforms annotated via pairwise genome alignments to *S. lucioperca* and *S. aurata* reference genomes. Species are grouped as Notothenioidei (blue), Cottioidei (orange), or other Perciformes (gray). Representative mutations across each exon for select species are shown to the right. Exons are shaded gray; deleted or missing exons are white with a red outline. Truncating variants are labeled by their type (e.g., frameshifts as +1 or –1), and splice site mutations are marked with red dashed lines at exon boundaries.

**Fig. S3.**
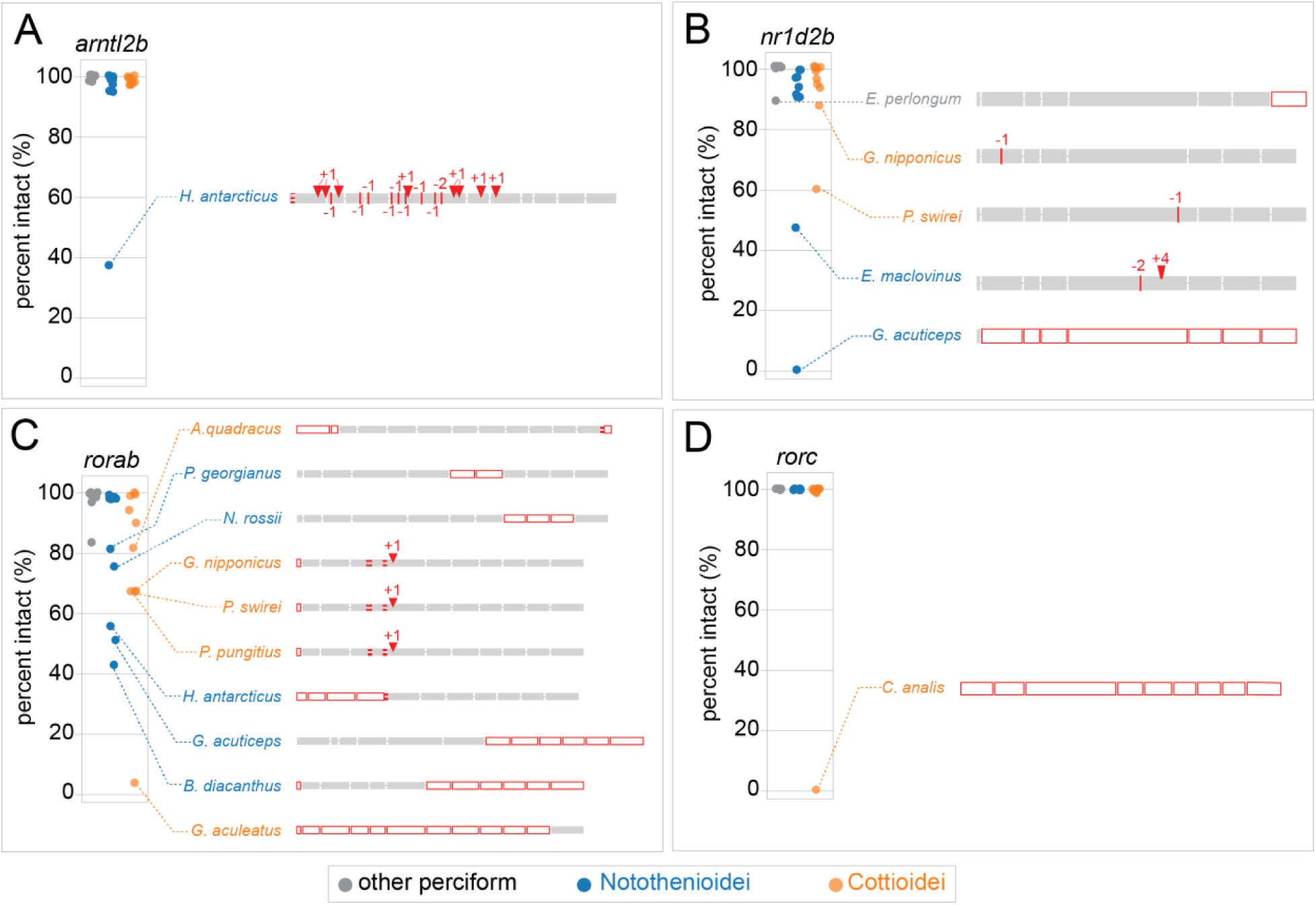
Example mutational variants across assorted biological clock genes. A) *arntl2b*, B) *nr1d2b*, C) *rorab*, D) *rorc*. Plots show the maximum percentage of intact coding sequence across all transcript isoforms annotated via pairwise genome alignments to *S. lucioperca* and *S. aurata* reference genomes. Species are grouped as Notothenioidei (blue), Cottioidei (orange), or other Perciformes (gray). Representative mutations across each exon for select species are shown to the right. Exons are shaded gray; deleted or missing exons are white with a red outline. Truncating variants are labeled by their type (e.g., frameshifts as +1 or –1), and splice site mutations are marked with red dashed lines at exon boundaries.

**Table S1.** Biological clock genes used in the analysis.

| **Gene name** | **Sander Ensembl ID** | **Sparus Ensembl ID** |
| --- | --- | --- |
| *arntl1a* | ENSSLUG00000025582 | ENSSAUG00010012423 |
| *arntl2a* | ENSSLUG00000025458 | ENSSAUG00010021136 |
| *arntl2b* | ENSSLUG00000023039 | ENSSAUG00010013586 |
| *clocka* | ENSSLUG00000011628 | ENSSAUG00010001441 |
| *clockb* | ENSSLUG00000011054 | ENSSAUG00010025019 |
| *cry-dash* | ENSSLUG00000012433 | ENSSAUG00010022353 |
| *cry1a* | ENSSLUG00000003675 | ENSSAUG00010012268 |
| *cry1b* | ENSSLUG00000002894 | ENSSAUG00010001264 |
| *cry2* | ENSSLUG00000025356 | ENSSAUG00010019549 |
| *cry3a* | ENSSLUG00000012954 | ENSSAUG00010004169 |
| *cry5* | ENSSLUG00000015034 | ENSSAUG00010015769 |
| *csnk1db* | ENSSLUG00000014906 | ENSSAUG00010004970 |
| *csnk1e* | ENSSLUG00000023267 | ENSSAUG00010009011 |
| *npas2* | ENSSLUG00000023767 | ENSSAUG00010018036 |
| *nr1d2a* | ENSSLUG00000011202 | ENSSAUG00010015973 |
| *nr1d2b* | ENSSLUG00000002539 | ENSSAUG00010013815 |
| *per1b* | ENSSLUG00000010714 | ENSSAUG00010014015 |
| *per2a* | ENSSLUG00000022765 | ENSSAUG00010010319 |
| *per2b* | ENSSLUG00000019172 | ENSSAUG00010027189 |
| *per3* | ENSSLUG00000011504 | ENSSAUG00010011519 |
| *roraa* | ENSSLUG00000015161 | ENSSAUG00010023974 |
| *rorab* | ENSSLUG00000016495 | ENSSAUG00010006455 |
| *rorb* | ENSSLUG00000022154 | ENSSAUG00010013250 |
| *rorc* | ENSSLUG00000010667 | ENSSAUG00010018260 |
| *rorca* | ENSSLUG00000007521 | ENSSAUG00010020966 |
| *rorcb* | ENSSLUG00000010091 | ENSSAUG00010018584 |
| *timeless* | ENSSLUG00000000306 | ENSSAUG00010024458 |

**Table S2.** Genome assemblies used in the analysis.

| Order | Suborder | Family | Species | GenBank Accession | Length (Mb) | Scaffold count | Scaffold N50 (Mb) | Contig count | Contig N50 (Mb) |
| --- | --- | --- | --- | --- | --- | --- | --- | --- | --- |
| Perciformes | Cottioidei | Anoplopomatidae | Anoplopoma fimbria | GCA_027596085.2 | 653.5 | 7493 | 26.7 | 8212 | 2.6 |
| Perciformes | Cottioidei | Cottidae | Clinocottus analis | GCA_023055335.1 | 538.1 | 443 | 21.0 | 662 | 9.2 |
| Perciformes | Cottioidei | Cyclopteridae | Cyclopterus lumpus | GCA_009769545.1 | 572.9 | 49 | 23.9 | 396 | 5.0 |
| Perciformes | Cottioidei | Liparidae | Pseudoliparis swirei | GCA_029220125.1 | 626.4 | 199 | 25.7 | 1102 | 4.2 |
| Perciformes | Cottioidei | Gasterosteidae | Apeltes quadracus | GCA_048569185.1 | 475.9 | 215 | 18.5 | 339 | 10.5 |
| Perciformes | Cottioidei | Gasterosteidae | Pungitius pungitius | GCA_949316345.1 | 480.5 | 175 | 21.0 | 913 | 1.4 |
| Perciformes | Cottioidei | Gasterosteidae | Gasterosteus nipponicus | GCA_014132575.2 | 599.5 | 3095 | 17.6 | 3718 | 0.5 |
| Perciformes | Cottioidei | Gasterosteidae | Gasterosteus aculeatus | GCA_016920845.1 | 471.9 | 2937 | 20.5 | 6059 | 0.5 |
| Perciformes | Cottioidei | Stichaeidae | Cebidichthys violaceus | GCA_008087265.1 | 593.0 | 467 | 6.7 | 505 | 5.5 |
| Perciformes | Cottioidei | Zoarcidae | Lycodopsis pacificus | GCA_028022725.1 | 646.4 | 109 | 27.7 | 475 | 4.7 |
| Perciformes | Notothenoidei | Artedidraconidae | Pogonophryne alpinna | GCA_028583405.1 | 1074.5 | 1111 | 41.8 | 3897 | 1.0 |
| Perciformes | Notothenoidei | Bathdraconidae | Gymnodraco acuticeps | GCA_902827175.1 | 996.9 | 2618 | 1.9 | 4183 | 0.5 |
| Perciformes | Notothenoidei | Bovichtidae | Cottoperca trigloides | GCA_900634415.1 | 609.4 | 322 | 25.2 | 766 | 6.3 |
| Perciformes | Notothenoidei | Bovichtidae | Bovichtus diacanthus | GCA_943590825.1 | 641.8 | 12443 | 7.3 | 44874 | 0.0 |
| Perciformes | Notothenoidei | Channichthyidae | Pseudochaenichthys georgianus | GCA_902827115.2 | 1026.2 | 1562 | 42.8 | 4129 | 0.7 |
| Perciformes | Notothenoidei | Channichthyidae | Chaenoccephalus aceratus | GCA_023974075.1 | 1065.6 | 3134 | 33.5 | 3856 | 1.5 |
| Perciformes | Notothenoidei | Eleginopidae | Eleginops maclovinus | GCA_036324505.1 | 606.3 | 26 | 26.7 | 406 | 7.6 |
| Perciformes | Notothenoidei | Harpagiferidae | Harpagifer antarcticus | GCA_902827135.1 | 941.5 | 1541 | 5.0 | 2579 | 1.1 |
| Perciformes | Notothenoidei | Nototheniidae | Trematomus bernacchii | GCA_902827165.1 | 867.1 | 864 | 8.8 | 1793 | 1.4 |
| Perciformes | Notothenoidei | Nototheniidae | Notothenia rossii | GCA_949606895.1 | 1042.9 | 943 | 89.7 | 5100 | 0.4 |
| Perciformes | Percoidei | Percidae | Gymnocephalus cernua | GCA_023631565.1 | 904.3 | 156 | 38.2 | 583 | 10.9 |
| Perciformes | Percoidei | Percidae | Sander lucioperca | GCA_008315115.1 | 900.5 | 1312 | 4.9 | 1347 | 4.7 |
| Spariformes | N/A | Sparidae | Sparus aurata | GCA_900880675.1 | 833.2 | 175 | 35.8 | 1223 | 2.9 |
| Perciformes | Scorpaenoidei | Scorpaenidae | Pterois miles | GCA_947000775.1 | 902.4 | N/A | N/A | 660 | 14.5 |
| Perciformes | Percoidei | Percidae | Perca flavescens | GCA_004354835.1 | 877.4 | 267 | 37.4 | 1096 | 4.3 |
| Perciformes | Percoidei | Percidae | Etheostoma perlongum | GCA_026937815.1 | 788.7 | 1179 | 30.8 | 2095 | 2.3 |
| Perciformes | Scorpaenoidei | Sebastidae | Sebastes schlegelii | GCA_014673565.1 | 848.0 | 1663 | 7.9 | 2110 | 5.5 |
| Perciformes | Scorpaenoidei | Synanceiidae | Synanceia verrucosa | GCA_029721515.1 | 691.9 | 1532 | 27.1 | 2325 | 12.0 |
| Perciformes | Serranoidei | Serranidae | Epinephelus lanceolatus | GCA_041903045.1 | 1089.4 | 176 | 45.8 | 183 | 44.4 |
| Perciformes | Serranoidei | Serranidae | Hypoplectrus puella | GCA_964304535.1 | 686.8 | 253 | 26.1 | 268 | 23.2 |
| Perciformes | Trigloidei | Triglidae | Chelidonichthys spinosus | GCA_029853015.1 | 624.7 | 293 | 28.1 | 406 | 13.8 |
| Perciformes | Percoidei | Trachinidae | Echiichthys vipera | GCA_963691815.1 | 800.4 | 133 | 33.0 | 739 | 2.6 |
| Perciformes | Serranoidei | Serranidae | Epinephelus cyanopodus | GCA_026686955.1 | 998.8 | 106 | 42.0 | 458 | 5.9 |
| Perciformes | Serranoidei | Serranidae | Centropristis striata | GCA_030273125.1 | 926.0 | 92 | 39.0 | 433 | 9.5 |
| Scombriformes | N/A | Scombridae | Thunnus maccoyii | GCA_910596095.1 | 782.4 | 57 | 33.8 | 152 | 26.8 |
| Pleuronectiformes | N/A | Pleuronectidae | Hippoglossus hippoglossus | GCA_009819705.1 | 596.8 | 56 | 26.3 | 346 | 7.0 |
| Cichliformes | N/A | Cichlidae | Oreochromis aureus | GCA_013358895.1 | 1000.0 | 303 | 40.7 | 1025 | 4.2 |
| Atheriniformes | N/A | Melanotaeniidae | Melanotaenia boesemani | GCA_017639745.1 | 865.6 | 92 | 37.9 | 532 | 9.3 |
| Acanthuriformes | N/A | Moronidae | Dicentrarchus labrax | GCA_905237075.1 | 695.9 | 302 | 29.9 | 574 | 12.7 |

**Table S3.** Relaxed selection across biological clock genes.

| Foreground branches | gene | Reference genome |  |  |  |
| --- | --- | --- | --- | --- | --- |
|  |  | S. lucioperca |  | S. auratus |  |
|  |  | K | p.adj | K | p.adj |
| Notothenioidei + Cottioidei | *arntl2a* | 0.82 | 3.76E-02 | 0.56 | 1.59E-02 |
| Notothenioidei + Cottioidei | *clocka* | 0.16 | 2.75E-03 | 0.72 | 5.01E-07 |
| Notothenioidei + Cottioidei | *cry1a* | 0.47 | 3.11E-06 | 0.45 | 2.25E-07 |
| Notothenioidei + Cottioidei | *cry5* | 0.85 | 2.17E-02 | 0.87 | 3.64E-02 |
| Notothenioidei + Cottioidei | *per1b* | 0.82 | 5.29E-08 | 0.00 | 4.74E-08 |
| Notothenioidei + Cottioidei | *per2a* | 0.51 | 6.15E-08 | 0.36 | 7.40E-04 |
| Notothenioidei + Cottioidei | *rorb* | 0.53 | 4.30E-05 | 0.50 | 1.80E-02 |
| Notothenioidei + Cottioidei | *rorc* | 0.79 | 1.71E-03 | 0.70 | 8.73E-05 |
| Notothenioidei + Cottioidei | *rorca* | 0.00 | 9.03E-03 | 0.25 | 2.47E-03 |
| Notothenioidei | *arntl2b* | 0.63 | 7.80E-02 | 0.50 | 2.47E-02 |
| Notothenioidei | *npas2* | 15.78 | 8.28E-02 | 8.40 | 2.51E-02 |
| Notothenioidei | *rorc* | 0.78 | 3.74E-02 | 0.72 | 2.80E-03 |
| Cottioidei | *cry1b* | 1.19 | 4.11E-02 | 1.16 | 6.77E-02 |
| Cottioidei | *clocka* | 0.70 | 1.52E-03 | 0.79 | 4.12E-04 |
| Cottioidei | *cry3a* | 0.71 | 1.51E-08 | 0.68 | 3.77E-05 |
| Cottioidei | *rorab* | 0.67 | 1.64E-03 | 0.83 | 5.27E-02 |
| Cottioidei | *cry1a* | 0.43 | 2.33E-05 | 0.69 | 1.07E-04 |
| Cottioidei | *rorca* | 0.63 | 6.35E-04 | 0.47 | 4.12E-04 |
| Cottioidei | *arntl2a* | 0.36 | 2.33E-05 | 0.14 | 1.09E-05 |

**Table S4.**
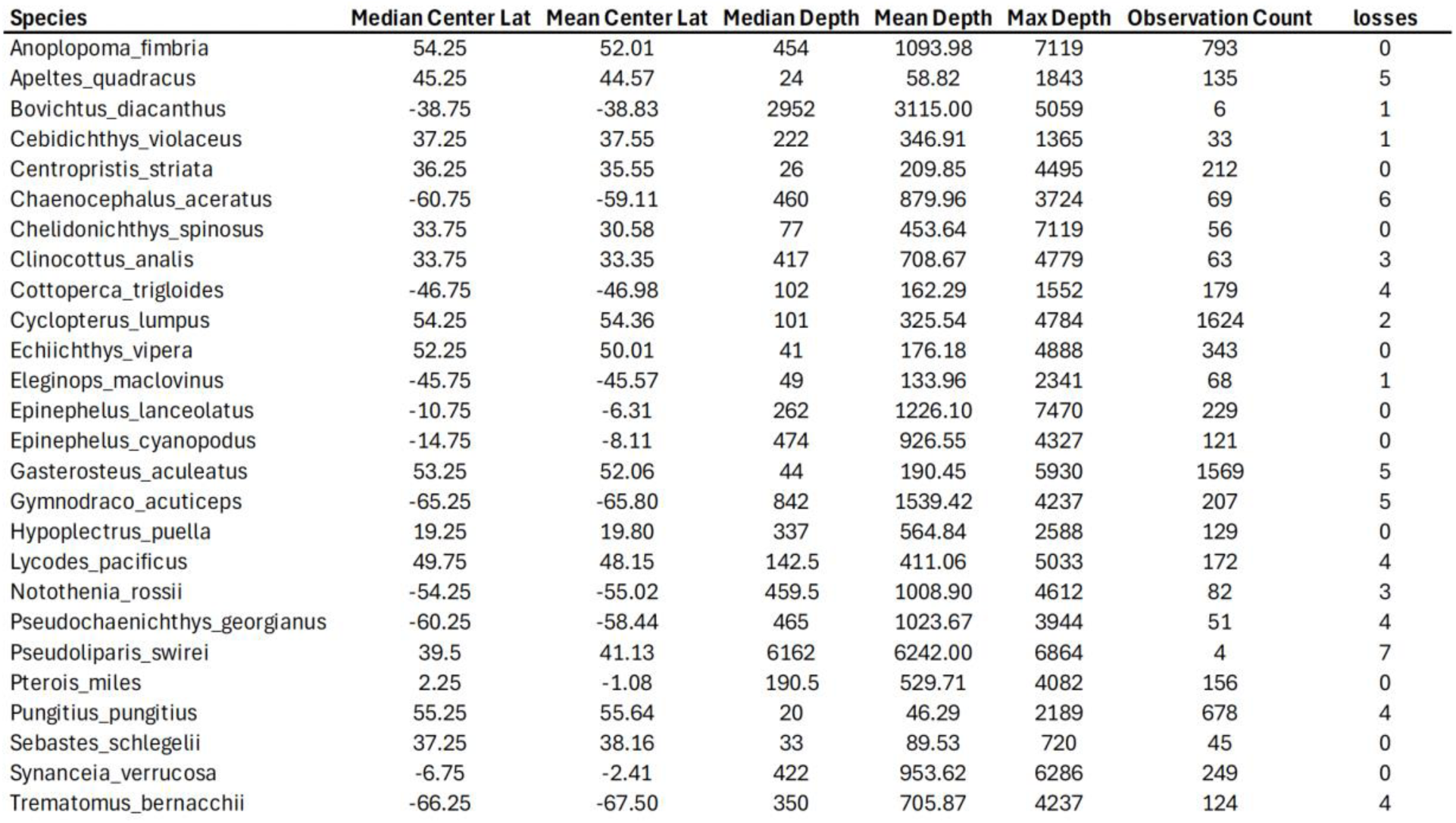
Species location data. Latitude and depth data from Aquamaps (88).

| Species | Median Center Lat | Mean Center Lat | Median Depth | Mean Depth | Max Depth | Observation Count | losses |
| --- | --- | --- | --- | --- | --- | --- | --- |
| Anoplopoma_fimbria | 54.25 | 52.01 | 454 | 1093.98 | 7119 | 793 | 0 |
| Apeltes_quadracus | 45.25 | 44.57 | 24 | 58.82 | 1843 | 135 | 5 |
| Bovichtus_diacanthus | -38.75 | -38.83 | 2952 | 3115.00 | 5059 | 6 | 1 |
| Cebidichthys_violaceus | 37.25 | 37.55 | 222 | 346.91 | 1365 | 33 | 1 |
| Centropristis_striata | 36.25 | 35.55 | 26 | 209.85 | 4495 | 212 | 0 |
| Chaenocephalus_aceratus | -60.75 | -59.11 | 460 | 879.96 | 3724 | 69 | 6 |
| Chelidonichthys_spinosus | 33.75 | 30.58 | 77 | 453.64 | 7119 | 56 | 0 |
| Clinocottus_analis | 33.75 | 33.35 | 417 | 708.67 | 4779 | 63 | 3 |
| Cottoperca_trigloides | -46.75 | -46.98 | 102 | 162.29 | 1552 | 179 | 4 |
| Cyclopterus_lumpus | 54.25 | 54.36 | 101 | 325.54 | 4784 | 1624 | 2 |
| Echiichthys_vipera | 52.25 | 50.01 | 41 | 176.18 | 4888 | 343 | 0 |
| Eleginops_maclovinus | -45.75 | -45.57 | 49 | 133.96 | 2341 | 68 | 1 |
| Epinephelus_lanceolatus | -10.75 | -6.31 | 262 | 1226.10 | 7470 | 229 | 0 |
| Epinephelus_cyanopodus | -14.75 | -8.11 | 474 | 926.55 | 4327 | 121 | 0 |
| Gasterosteus_aculeatus | 53.25 | 52.06 | 44 | 190.45 | 5930 | 1569 | 5 |
| Gymnodraco_acuticeps | -65.25 | -65.80 | 842 | 1539.42 | 4237 | 207 | 5 |
| Hypoplectrus_puella | 19.25 | 19.80 | 337 | 564.84 | 2588 | 129 | 0 |
| Lycodes_pacificus | 49.75 | 48.15 | 142.5 | 411.06 | 5033 | 172 | 4 |
| Notothenia_rossii | -54.25 | -55.02 | 459.5 | 1008.90 | 4612 | 82 | 3 |
| Pseudochaenichthys_georgianus | -60.25 | -58.44 | 465 | 1023.67 | 3944 | 51 | 4 |
| Pseudoliparis_swirei | 39.5 | 41.13 | 6162 | 6242.00 | 6864 | 4 | 7 |
| Pterois_miles | 2.25 | -1.08 | 190.5 | 529.71 | 4082 | 156 | 0 |
| Pungitius_pungitius | 55.25 | 55.64 | 20 | 46.29 | 2189 | 678 | 4 |
| Sebastes_schlegelii | 37.25 | 38.16 | 33 | 89.53 | 720 | 45 | 0 |
| Synanceia_verrucosa | -6.75 | -2.41 | 422 | 953.62 | 6286 | 249 | 0 |
| Trematomus_bernacchii | -66.25 | -67.50 | 350 | 705.87 | 4237 | 124 | 4 |

**Table S5.** Breakdown of global gene status (TOGA) by reference genome.

| Reference | Species | I+PI | L+UL | I | PI | L | UL | M | PM | PG |
| --- | --- | --- | --- | --- | --- | --- | --- | --- | --- | --- |
| sparus_aurata | anoplopoma_fimbria | 72.3 | 26.5 | 71.2 | 1.0 | 11.6 | 14.9 | 0.7 | 0.3 | 0.3 |
| sander_luciperca | anoplopoma_fimbria | 69.1 | 27.3 | 68.3 | 0.8 | 9.7 | 17.6 | 2.0 | 0.4 | 1.2 |
| merged | anoplopoma_fimbria | 76.1 | 20.7 | 75.3 | 0.8 | 9.5 | 11.2 | 1.9 | 0.3 | 1.0 |
| sparus_aurata | apeltes_quadracus | 62.8 | 34.6 | 61.9 | 0.9 | 18.0 | 16.6 | 1.5 | 0.3 | 0.8 |
| sander_luciperca | apeltes_quadracus | 59.6 | 32.3 | 59.0 | 0.5 | 13.9 | 18.4 | 3.9 | 0.4 | 3.8 |
| merged | apeltes_quadracus | 66.7 | 26.8 | 66.1 | 0.6 | 14.1 | 12.7 | 3.7 | 0.4 | 2.4 |
| sparus_aurata | bovichtus_diacanthus | 67.6 | 21.9 | 60.8 | 6.8 | 12.8 | 9.1 | 5.9 | 1.8 | 2.8 |
| sander_luciperca | bovichtus_diacanthus | 66.7 | 20.0 | 60.2 | 6.4 | 8.2 | 11.9 | 7.9 | 1.8 | 3.3 |
| merged | bovichtus_diacanthus | 73.0 | 15.4 | 66.1 | 6.9 | 8.7 | 6.7 | 7.4 | 1.5 | 2.7 |
| sparus_aurata | cebidichthys_violaceus | 72.7 | 23.3 | 71.0 | 1.7 | 13.1 | 10.3 | 2.4 | 0.4 | 1.1 |
| sander_luciperca | cebidichthys_violaceus | 68.5 | 24.1 | 67.2 | 1.3 | 10.3 | 13.8 | 4.3 | 0.5 | 2.6 |
| merged | cebidichthys_violaceus | 75.4 | 18.7 | 74.0 | 1.3 | 10.9 | 7.8 | 3.6 | 0.5 | 1.9 |
| sparus_aurata | centropristis_striata | 79.8 | 19.3 | 78.8 | 1.0 | 9.7 | 9.6 | 0.5 | 0.2 | 0.2 |
| sander_luciperca | centropristis_striata | 74.7 | 22.1 | 73.8 | 0.9 | 8.4 | 13.7 | 1.6 | 0.4 | 1.3 |
| merged | centropristis_striata | 82.2 | 15.1 | 81.3 | 0.9 | 7.9 | 7.2 | 1.4 | 0.3 | 1.0 |
| sparus_aurata | chaenocephalus_aceratus | 53.0 | 40.0 | 51.6 | 1.5 | 18.9 | 21.2 | 3.5 | 0.9 | 2.5 |
| sander_luciperca | chaenocephalus_aceratus | 51.8 | 39.3 | 50.4 | 1.3 | 16.4 | 23.0 | 4.8 | 1.0 | 3.2 |
| merged | chaenocephalus_aceratus | 57.2 | 35.4 | 55.7 | 1.5 | 17.8 | 17.6 | 4.5 | 0.8 | 2.2 |
| sparus_aurata | chelidonichthys_spinosus | 70.3 | 28.7 | 69.5 | 0.9 | 13.6 | 15.1 | 0.4 | 0.4 | 0.1 |
| sander_luciperca | chelidonichthys_spinosus | 66.5 | 28.6 | 65.8 | 0.7 | 11.5 | 17.1 | 2.6 | 0.5 | 1.9 |
| merged | chelidonichthys_spinosus | 73.6 | 22.5 | 72.9 | 0.7 | 11.4 | 11.1 | 2.3 | 0.4 | 1.3 |
| sparus_aurata | clinocottus_analis | 74.2 | 23.9 | 73.3 | 0.9 | 14.2 | 9.7 | 1.0 | 0.3 | 0.7 |
| sander_luciperca | clinocottus_analis | 70.2 | 23.8 | 69.6 | 0.7 | 11.0 | 12.9 | 2.9 | 0.4 | 2.6 |
| merged | clinocottus_analis | 77.7 | 17.6 | 77.0 | 0.7 | 10.8 | 6.8 | 2.6 | 0.4 | 1.8 |
| sparus_aurata | cottoperca_gobio | 64.1 | 31.4 | 63.0 | 1.1 | 15.3 | 16.2 | 2.9 | 0.5 | 1.1 |
| sander_luciperca | cottoperca_gobio | 62.6 | 31.0 | 61.6 | 1.0 | 12.3 | 18.7 | 4.0 | 0.7 | 1.7 |
| merged | cottoperca_gobio | 68.8 | 25.7 | 67.7 | 1.1 | 12.8 | 12.9 | 3.6 | 0.6 | 1.3 |
| sparus_aurata | cyclopterus_lumpus | 70.0 | 27.8 | 69.1 | 0.8 | 16.1 | 11.8 | 1.4 | 0.3 | 0.5 |
| sander_luciperca | cyclopterus_lumpus | 66.7 | 27.0 | 66.0 | 0.8 | 12.5 | 14.5 | 3.7 | 0.5 | 2.1 |
| merged | cyclopterus_lumpus | 73.9 | 20.9 | 73.1 | 0.8 | 12.6 | 8.4 | 3.3 | 0.4 | 1.5 |
| sparus_aurata | dicentrarchus_labrax | 81.5 | 18.0 | 80.4 | 1.1 | 7.9 | 10.1 | 0.3 | 0.2 | 0.1 |
| sander_luciperca | dicentrarchus_labrax | 74.4 | 22.7 | 73.5 | 0.8 | 8.5 | 14.2 | 1.4 | 0.4 | 1.2 |
| merged | dicentrarchus_labrax | 82.0 | 15.4 | 81.2 | 0.8 | 7.8 | 7.5 | 1.3 | 0.3 | 1.0 |
| sparus_aurata | echiichthys_vipera | 76.2 | 22.8 | 75.0 | 1.2 | 12.8 | 10.1 | 0.4 | 0.4 | 0.2 |
| sander_luciperca | echiichthys_vipera | 72.8 | 23.3 | 71.8 | 1.0 | 9.7 | 13.6 | 1.9 | 0.4 | 1.7 |
| merged | echiichthys_vipera | 80.0 | 16.5 | 79.0 | 1.0 | 9.3 | 7.2 | 1.8 | 0.4 | 1.3 |
| sparus_aurata | eleginops_maclovinus | 68.0 | 29.7 | 67.2 | 0.8 | 15.0 | 14.7 | 1.3 | 0.4 | 0.6 |
| sander_lucioperca | eleginops_maclovinus | 66.0 | 28.6 | 65.2 | 0.8 | 11.5 | 17.2 | 2.7 | 0.5 | 2.1 |
| merged | eleginops_maclovinus | 66.7 | 30.9 | 65.8 | 0.9 | 9.4 | 21.6 | 1.3 | 0.2 | 0.8 |
| sander_lucioperca | epinephalus_cyanopodus | 66.7 | 31.0 | 65.8 | 0.9 | 9.4 | 21.6 | 1.3 | 0.2 | 0.8 |
| sparus_aurata | epinephalus_cyanopodus | 71.1 | 28.3 | 70.2 | 0.9 | 9.7 | 18.6 | 0.3 | 0.2 | 0.1 |
| merged | epinephalus_cyanopodus | 73.3 | 24.6 | 72.4 | 0.9 | 9.4 | 15.3 | 1.2 | 0.2 | 0.7 |
| sparus_aurata | epinephelus_lanceolatus | 78.4 | 19.7 | 75.9 | 2.5 | 8.9 | 10.7 | 1.4 | 0.3 | 0.1 |
| sander_lucioperca | epinephelus_lanceolatus | 74.1 | 21.3 | 72.1 | 2.0 | 7.5 | 13.7 | 3.2 | 0.5 | 1.0 |
| merged | epinephelus_lanceolatus | 81.1 | 14.7 | 79.0 | 2.0 | 7.2 | 7.5 | 3.0 | 0.4 | 0.8 |
| sparus_aurata | etheostoma_perlongum | 58.4 | 38.9 | 57.3 | 1.1 | 14.4 | 24.5 | 1.9 | 0.4 | 0.5 |
| sander_lucioperca | etheostoma_perlongum | 62.5 | 35.5 | 61.3 | 1.3 | 9.1 | 26.5 | 1.2 | 0.4 | 0.3 |
| merged | etheostoma_perlongum | 67.5 | 30.8 | 66.2 | 1.3 | 9.6 | 21.2 | 1.2 | 0.3 | 0.3 |
| sparus_aurata | gasterosteus_aculeatus | 65.5 | 29.1 | 64.3 | 1.3 | 16.0 | 13.1 | 4.1 | 0.4 | 0.9 |
| sander_lucioperca | gasterosteus_aculeatus | 62.9 | 27.4 | 61.9 | 1.0 | 12.2 | 15.2 | 5.5 | 0.6 | 3.5 |
| merged | gasterosteus_aculeatus | 70.1 | 21.9 | 69.0 | 1.1 | 12.3 | 9.6 | 5.3 | 0.5 | 2.2 |
| sparus_aurata | gasterosteus_nipponicus | 56.9 | 39.6 | 55.5 | 1.4 | 17.7 | 21.8 | 2.0 | 0.4 | 1.2 |
| sander_lucioperca | gasterosteus_nipponicus | 54.6 | 36.9 | 53.5 | 1.1 | 14.1 | 22.8 | 4.1 | 0.6 | 3.8 |
| merged | gasterosteus_nipponicus | 61.3 | 31.8 | 60.2 | 1.2 | 14.4 | 17.4 | 4.0 | 0.5 | 2.4 |
| sparus_aurata | gymnocephalus_cernua | 78.3 | 20.6 | 77.1 | 1.2 | 10.6 | 10.1 | 0.6 | 0.3 | 0.2 |
| sander_lucioperca | gymnocephalus_cernua | 83.2 | 16.0 | 82.1 | 1.1 | 4.6 | 11.4 | 0.5 | 0.2 | 0.1 |
| merged | gymnocephalus_cernua | 88.0 | 11.4 | 86.8 | 1.2 | 4.5 | 6.8 | 0.4 | 0.2 | 0.1 |
| sparus_aurata | gymnodraco_acuticeps | 63.4 | 29.1 | 61.2 | 2.2 | 14.1 | 15.0 | 3.3 | 1.0 | 3.2 |
| sander_lucioperca | gymnodraco_acuticeps | 61.5 | 28.7 | 59.6 | 1.9 | 11.4 | 17.3 | 5.0 | 1.3 | 3.6 |
| merged | gymnodraco_acuticeps | 67.9 | 23.7 | 65.7 | 2.2 | 12.0 | 11.6 | 4.7 | 1.1 | 2.7 |
| sparus_aurata | hippoglossus_hippoglossus | 71.5 | 25.3 | 70.8 | 0.7 | 15.2 | 10.2 | 2.1 | 0.3 | 0.7 |
| sander_lucioperca | hippoglossus_hippoglossus | 67.3 | 25.5 | 66.8 | 0.5 | 12.2 | 13.3 | 4.4 | 0.5 | 2.4 |
| merged | hippoglossus_hippoglossus | 74.9 | 18.9 | 74.4 | 0.5 | 11.9 | 7.0 | 4.1 | 0.4 | 1.7 |
| sparus_aurata | hypoplectrus_puella | 74.7 | 20.5 | 66.9 | 7.8 | 10.5 | 9.9 | 3.5 | 0.4 | 0.9 |
| sander_lucioperca | hypoplectrus_puella | 70.4 | 21.3 | 64.1 | 6.3 | 8.6 | 12.7 | 4.7 | 0.6 | 3.1 |
| merged | hypoplectrus_puella | 77.7 | 15.1 | 70.7 | 7.0 | 8.3 | 6.8 | 4.5 | 0.6 | 2.1 |
| sparus_aurata | lycodes_pacificus | 76.9 | 21.7 | 75.8 | 1.1 | 12.0 | 9.7 | 0.9 | 0.2 | 0.3 |
| sander_lucioperca | lycodes_pacificus | 72.1 | 23.8 | 71.2 | 0.9 | 10.2 | 13.7 | 2.2 | 0.4 | 1.5 |
| merged | lycodes_pacificus | 79.4 | 17.0 | 78.5 | 0.9 | 9.8 | 7.2 | 2.1 | 0.3 | 1.2 |
| sparus_aurata | melanotaenia_boesmani | 73.1 | 24.2 | 72.2 | 0.8 | 14.3 | 9.9 | 1.6 | 0.3 | 0.9 |
| sander_lucioperca | melanotaenia_boesmani | 66.8 | 25.1 | 66.3 | 0.6 | 12.4 | 12.7 | 3.5 | 0.6 | 4.0 |
| merged | melanotaenia_boesmani | 75.4 | 18.6 | 74.7 | 0.7 | 12.0 | 6.6 | 3.3 | 0.5 | 2.3 |
| sparus_aurata | notothenia_rossii | 73.5 | 23.6 | 69.7 | 3.8 | 12.3 | 11.2 | 2.2 | 0.3 | 0.5 |
| sander_lucioperca | notothenia_rossii | 69.3 | 25.0 | 65.9 | 3.4 | 10.3 | 14.7 | 3.6 | 0.4 | 1.6 |
| merged | notothenia_rossii | 76.6 | 18.9 | 72.8 | 3.7 | 10.5 | 8.5 | 3.0 | 0.4 | 1.1 |
| sparus_aurata | oreochromis_aureus | 75.6 | 21.7 | 74.7 | 0.9 | 12.0 | 9.7 | 1.5 | 0.3 | 0.9 |
| sander_lucioperca | oreochromis_aureus | 68.8 | 23.5 | 68.1 | 0.7 | 10.5 | 12.9 | 3.7 | 0.5 | 3.6 |
| merged | oreochromis_aureus | 77.4 | 16.5 | 76.7 | 0.7 | 10.1 | 6.4 | 3.5 | 0.4 | 2.2 |
| sparus_aurata | perca_flavescens | 75.9 | 22.5 | 74.8 | 1.0 | 10.9 | 11.7 | 1.0 | 0.3 | 0.3 |
| sander_luciperca | perca_flavescens | 82.5 | 17.0 | 81.5 | 1.1 | 4.4 | 12.6 | 0.2 | 0.2 | 0.1 |
| merged | perca_flavescens | 87.1 | 12.4 | 86.0 | 1.1 | 4.4 | 8.0 | 0.2 | 0.2 | 0.1 |
| sparus_aurata | pogonophryne_albipinna | 73.0 | 24.2 | 71.3 | 1.7 | 12.9 | 11.2 | 1.6 | 0.4 | 0.8 |
| sander_luciperca | pogonophryne_albipinna | 69.3 | 25.3 | 67.8 | 1.5 | 10.9 | 14.4 | 2.9 | 0.5 | 1.9 |
| merged | pogonophryne_albipinna | 76.4 | 19.2 | 74.8 | 1.6 | 11.2 | 8.0 | 2.7 | 0.4 | 1.3 |
| sparus_aurata | pseudochaenichthys_georgianus | 66.0 | 26.5 | 64.1 | 1.9 | 14.3 | 12.2 | 4.3 | 0.9 | 2.2 |
| sander_luciperca | pseudochaenichthys_georgianus | 64.0 | 26.5 | 62.3 | 1.7 | 11.6 | 14.9 | 5.3 | 1.0 | 3.2 |
| merged | pseudochaenichthys_georgianus | 70.4 | 21.5 | 68.5 | 1.9 | 12.4 | 9.1 | 5.1 | 0.7 | 2.2 |
| sparus_aurata | pseudoliparis_swirei | 63.8 | 31.6 | 62.8 | 0.9 | 20.4 | 11.2 | 3.1 | 0.4 | 1.0 |
| sander_luciperca | pseudoliparis_swirei | 61.4 | 28.3 | 60.6 | 0.8 | 14.5 | 13.9 | 6.2 | 0.7 | 3.5 |
| merged | pseudoliparis_swirei | 68.5 | 23.5 | 67.7 | 0.8 | 15.5 | 8.0 | 5.4 | 0.5 | 2.0 |
| sparus_aurata | pterois_miles | 73.9 | 24.8 | 73.0 | 0.9 | 12.3 | 12.5 | 0.7 | 0.3 | 0.3 |
| sander_luciperca | pterois_miles | 70.6 | 26.0 | 69.9 | 0.7 | 10.2 | 15.8 | 1.8 | 0.3 | 1.2 |
| merged | pterois_miles | 77.7 | 19.2 | 77.0 | 0.7 | 9.8 | 9.5 | 1.7 | 0.3 | 1.0 |
| sparus_aurata | pungitius_pungitius | 71.6 | 25.8 | 70.3 | 1.3 | 15.7 | 10.1 | 1.7 | 0.3 | 0.7 |
| sander_luciperca | pungitius_pungitius | 67.1 | 25.4 | 66.1 | 1.0 | 12.3 | 13.1 | 3.7 | 0.5 | 3.3 |
| merged | pungitius_pungitius | 75.2 | 19.0 | 74.2 | 1.1 | 12.2 | 6.8 | 3.4 | 0.4 | 2.1 |
| sparus_aurata | sander_luciperca | 73.1 | 25.4 | 72.0 | 1.1 | 10.7 | 14.7 | 0.7 | 0.4 | 0.5 |
| sparus_aurata | sebastes_schlegelii | 74.9 | 23.2 | 73.8 | 1.1 | 10.4 | 12.8 | 1.3 | 0.3 | 0.3 |
| sander_luciperca | sebastes_schlegelii | 70.8 | 25.4 | 69.9 | 0.9 | 9.3 | 16.1 | 2.4 | 0.4 | 1.1 |
| merged | sebastes_schlegelii | 77.8 | 18.8 | 76.8 | 1.0 | 9.1 | 9.7 | 2.2 | 0.3 | 0.9 |
| sparus_aurata | synanceia_verrucosa | 74.6 | 23.8 | 73.8 | 0.9 | 14.4 | 9.3 | 0.8 | 0.5 | 0.3 |
| sander_luciperca | synanceia_verrucosa | 70.5 | 23.4 | 70.0 | 0.5 | 10.9 | 12.5 | 3.3 | 0.5 | 2.3 |
| merged | synanceia_verrucosa | 77.8 | 17.1 | 77.3 | 0.5 | 10.9 | 6.2 | 3.0 | 0.5 | 1.6 |
| sparus_aurata | thunnus_maccoyii | 79.8 | 19.0 | 79.0 | 0.9 | 9.4 | 9.6 | 0.7 | 0.3 | 0.2 |
| sander_luciperca | thunnus_maccoyii | 74.1 | 22.3 | 73.4 | 0.8 | 9.0 | 13.3 | 2.0 | 0.3 | 1.3 |
| merged | thunnus_maccoyii | 81.7 | 15.0 | 81.0 | 0.7 | 8.3 | 6.7 | 1.9 | 0.3 | 1.1 |
| sparus_aurata | trematomus_bernacchii | 69.0 | 26.9 | 67.9 | 1.1 | 12.9 | 13.9 | 2.1 | 0.6 | 1.5 |
| sander_luciperca | trematomus_bernacchii | 66.2 | 27.7 | 65.1 | 1.1 | 11.0 | 16.7 | 3.4 | 0.6 | 2.1 |
| merged | trematomus_bernacchii | 73.0 | 21.6 | 71.9 | 1.1 | 11.1 | 10.5 | 3.3 | 0.5 | 1.6 |

